# Flexible analysis of animal behavior via time-resolved manifold embedding

**DOI:** 10.1101/2020.09.30.321406

**Authors:** Ryan A. York, Arnaldo Carreira-Rosario, Lisa M. Giocomo, Thomas R. Clandinin

## Abstract

Uncovering relationships between neural activity and behavior represents a critical challenge, one that would benefit from facile tools that can capture complex structures within large datasets. Here we demonstrate a generalizable strategy for capturing such structures across diverse behaviors: Time-REsolved BehavioraL Embedding (TREBLE). Using data from synthetic trajectories, adult and larval *Drosophila*, and mice we show how TREBLE captures both continuous and discrete behavioral dynamics, can uncover variation across individuals, detect the effects of optogenetic perturbation in unbiased fashion, and reveal structure in pose estimation data. By applying TREBLE to moving mice, and medial entorhinal cortex (MEC) recordings, we show that nearly all MEC neurons encode information relevant to specific movement patterns, expanding our understanding of how navigation is related to the execution of locomotion. Thus, TREBLE provides a flexible framework for describing the structure of complex behaviors and their relationships to neural activity.

## Main

Accurate descriptions of animal behavior are essential to understanding brain function. However, naturalistic behavior in freely moving animals is often continuous, has structure across multiple timescales, and can vary broadly between individuals and contexts. Thus, statistical tools that can capture and visualize the temporal structural of behavior, can operate on large data sets derived from many individuals, and are generalizable across systems and experiments are of central interest.

Advances in tracking technology have enabled measurements of animal movement from a wide range of species (Datta et al. 2019; Pereira et al. 2020; Mathis et al. 2020). These datasets are often extremely rich, and include correlated movements across timescales, features that must be accounted for in efforts to link neural activity to behavior. In parallel, a variety of sophisticated methods have emerged to parse such behavioral structure and to measure behavioral changes caused by ever more powerful experimental perturbations (Brown & de Bivort 2018; Datta et al. 2019; Pereira et al. 2020; Mathis et al. 2020). However, many of these methods are complex to implement, and require extensive adaptation to specific species, contexts and experimental goals. As a result, many investigators continue to rely upon either instantaneous measures of specific behavioral parameters (such as velocities), or standard dimensionality reduction approaches in which relatively little variance in the behavior is accounted for in the first two dimensions. We therefore reasoned that an accessible framework that could be easily applied to a wide variety of species and contexts, and which would allow the temporal structure of behavior to be embedded and visualized in a low dimensional manifold, would be of widespread utility.

Here we describe Time-REsolved BehavioraL Embedding (TREBLE), an easy-to-use method that can describe the structure of behavior, assess the effects of experimental perturbations across the entire space of behavioral measurements, and provide intuitive representations of the relationships between neural activity and behavior. We reasoned that individual movements, by analogy with some genomic analyses, could be treated akin to conserved nucleotide sequence blocks of variable length. In genomics, quantitative and qualitative differences in sequence can be efficiently revealed by shotgunning long sequences into smaller, overlapping blocks that preserve local structure (Staden 1979; Wang et al. 2009). These blocks are then assembled into libraries that densely sample the structure, frequency and surrounding sequence of each block, while smoothly reconstructing the entire sequence. Quantitative variation in these features can then be assessed using standardized computational methods often including dimensionality reduction followed by statistical testing. Building on this conceptual parallel, we developed TREBLE as a method for extracting all behavioral sequences from a dataset (analogous to shotgun sequencing), assembling ‘libraries’, and creating a shared ‘behavior space’ using dimensionality reduction.

In the TREBLE framework, behavior is first quantitatively measured (Figure 1A). Relevant measurements such as centroid velocities, or changes in body or limb position, are computed from video data, and segmented into highly overlapped temporal windows, the size of which is constrained by the temporal structure of behavior. These windows are then collected into a large library encompassing all individuals and experimental perturbations and assembled in a low-dimensional behavior space. Depending on window size, this space can flexibly capture a range of temporal dynamics, ranging from unique trajectories to recurring patterns of movement. The resulting space can be leveraged to decode complex patterns of movement in a time-resolved fashion, facilitating analyses such as the rapid comparison of individuals or identification of neural perturbations and stimulus effects.

**Figure 1.**
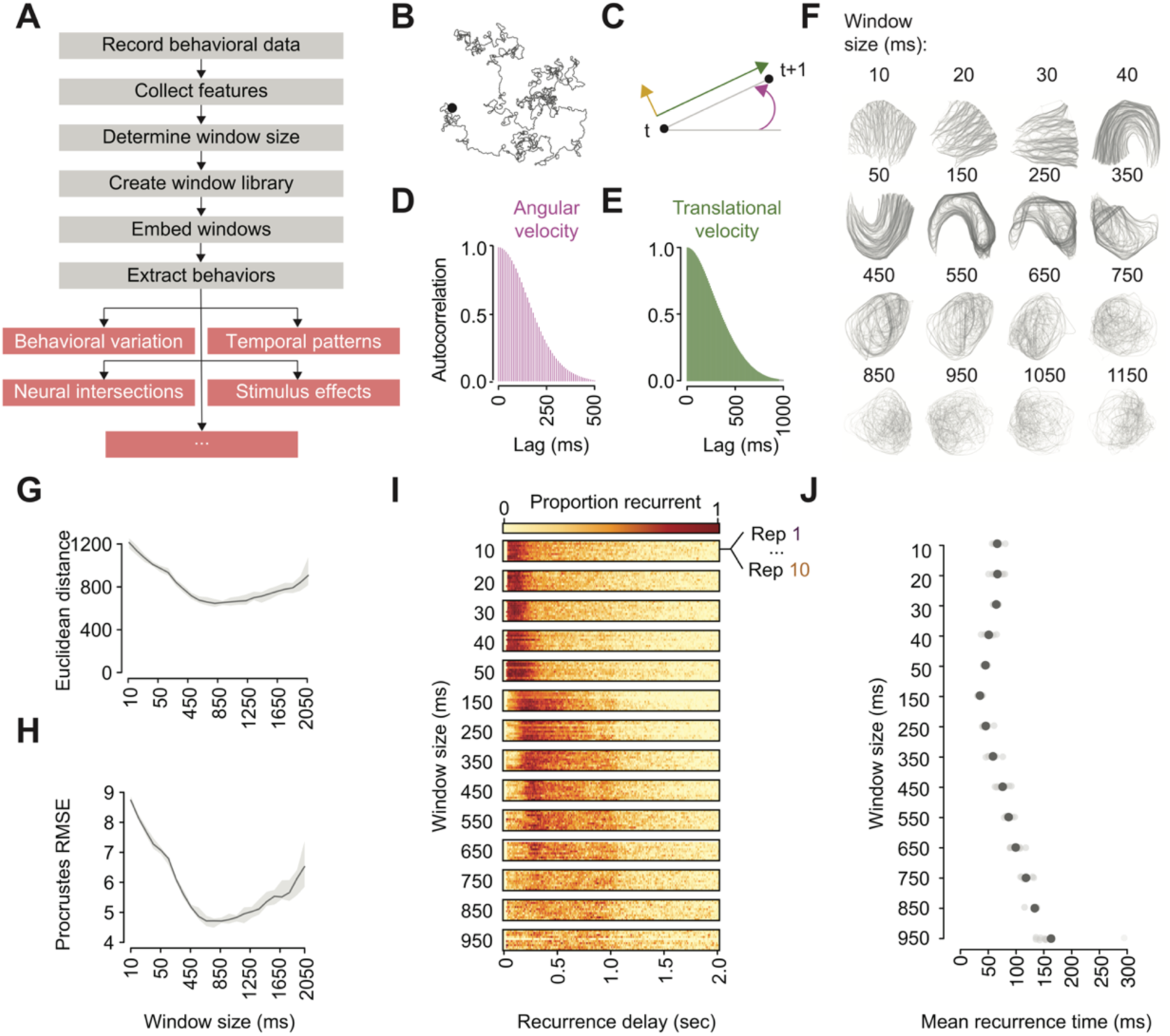
The TREBLE framework and its application to synthetic data. **(A)** Outline of the TREBLE framework. Behavioral data are recorded. Relevant features (such as velocities) are extracted these recordings following which empirical analyses are used to determine the optimal sampling window size. The chosen window size is then used to create a library spanning all time points in the data set. The resulting window library is embedded into a low-dimensional behavior space from which recurrent behaviors can be decoded and used for a number of analyses (examples labeled in red boxes). **(B)** Example correlated random walk used for parameter tuning (see also Figure S1). **(C)** Velocity components that can be calculated from movements in a 2-d plane. The black point at time *t* denotes the beginning of the trajectory which proceeds to *t+1*. The purple arrow corresponds to the angular velocity of this trajectory while the green and yellow arrows represent translational velocity and side slip, respectively. **(D)** The observed autocorrelation distribution of angular velocity computed from all correlated random walks. **(E)** The observed autocorrelation distribution of translational velocity. **(F)** Example behavior spaces for a range of window sizes. Spaces are plotted here by connecting temporally adjacent points (corresponding to feature windows), gray lines. Darker lines reflect repeated visits along the same pathway. **(G)** Mean intra-point Euclidean distances as a function of window size. Mean values (dark gray); Standard error of the mean (shading). **(H)** Procrustes distance RMSE measures as a function of window size. Mean values (dark gray); Standard error of the mean (light gray shading). **(I)** Recurrence plot of behavior spaces produced from the correlated random walk dataset. The proportion of recurrent points given a range of time delays spanning from 0 to 2 seconds, is indicated by the color of the corresponding bins (ranging from light yellow to dark red). Each window size contains the distributions for 10 replicate walks. **(J)** Mean recurrence times as a function of window size. Population mean (large dark circle); Individual replicates (small light circle).

Here, we use TREBLE to analyze a wide variety of data sources (synthetic trajectories, adult and larval fruit flies, mice (with both 2D and 3D pose tracking)) and applications (behavioral repertoire description, experimental perturbation, intersection of behavior with imaging and electrophysiological neuron recordings). We show that TREBLE’s parameters can be rationally chosen using empirical analyses and that the resulting behavior spaces are regularly structured and display recurrent dynamics. Individual movement bouts can be easily decoded and classified, the representation of which can be tailored to a user’s need and allowing for detailed dissections of behavior into its constituent components. TREBLE can handle data from many individuals and millions of data points simultaneously, making possible the detection of otherwise invisible behavioral variation. Finally, we demonstrate the ability of TREBLE to rapidly uncover neurobehavioral relationships during optogenetic perturbations and neural imaging in flies, as well as electrophysiological recordings in freely moving mice.

## Results

### Calibrating TREBLE

To calibrate this approach, we first used a large, synthetic dataset comprised of correlated random walks to explore the relationships between window size and the emergent structure of behavior space (Bovet 1988) (Figure 1B; Figures S1A-B). Individual random walks were completely described by a combination of velocity features (translational velocity, angular velocity, and side slip; Figure 1C) with characteristic temporal dependencies determined by the correlated random walk generator (and highlighted by their autocorrelations; Figures 1D-E). We reasoned that window sizes smaller than the underlying correlations within and between velocities would artificially granularize features, while windows larger than the underlying correlations would combine uncorrelated features. Given this, an ‘ideal’ window size should show less variance across trials and display recurrent dynamics, paths through the space that repeat independently in the dataset since stereotyped movements that repeat would necessarily lead to recurrent paths in such a space.

To explore this, we assessed the effect of window size on feature representation, sweeping sizes from 10ms (containing 1 window, reflecting an instantaneous measurement of features) to 2 seconds (see Methods). Windows extracted from all replicate walk trajectories were then embedded into behavior space using the UMAP algorithm, a computationally efficient, non-linear dimensionality reduction approach (McInnes et al. 2018). The structure of behavior space varied broadly as a function of window size; topologies ranged between disordered (10-30ms), recurrent (40∼350ms), and unique paths (450ms ∼ 2 seconds) (Figure 1F; Figure S1B). These observations confirmed our initial intuition that window size could have a range of effects on the structure of behavior space and that more ordered spaces could be derived by considering timescales longer than raw/instantaneous measurements.

We next sought to quantify this topological variation. We employed two statistics, one targeting local structural differences and the other focused on global variation. To assess local variation, we measured the average Euclidean distance between temporally adjacent points, reasoning that spaces with smoother paths would display smaller distances with less variance. Mean values of Euclidean distance displayed a trough spanning window sizes of between 450 to 1250ms (Figure 1G). The coefficient of variation (CV) highlighted a broad overlapping region of similarity, spanning window sizes of between 10 to 950ms (Figure S1C). To assess global variation, we measured Procrustes distance (McInnes et al. 2018; Dryden & Mardia 1998). This metric compares the difference between configurations of points, used here to quantify the distance between replicate behavior spaces of a given window size. This measure displayed a pattern similar to Euclidean distance, with a large trough between 450 and 1250ms, and a consistent CV up to window sizes of 950ms (Figure 1H; Figure S1D). Thus, these metrics allow quantification of the effects of window size, in this case revealing stability in topologies of behavior space across a range of windows.

We then explored how the choice of window size affects the temporal progression of trajectories through behavior space by measuring the frequency with which portions of a trajectory were repeated. We delineated a neighborhood around each point in behavior space (see Methods) and determined the amount of time between visits to each neighborhood. This was then represented as the proportion of all neighborhoods that were revisited (Bruno et al. 2017). The distribution of these values was consistent across replicates *within* a given window size but varied across sizes (Figure 1I). Window sizes of 150 to 650ms displayed a dominant return time of approximately 250ms, with a secondary peak at 1 second (Figure 1I). Windows larger than 650ms displayed increasingly noisy return time distributions (Figure 1I). Reflecting this, plots of mean recurrence times revealed a structured tuning curve in which the mean and variance were minimized at a window size of 150ms (Figure 1J). Combining these observations, the broad tuning curves in both the spatial and temporal properties of behavior space indicate that while window size can have appreciable effects, this representation of behavior can be robust across multiple parameter values.

### Analysis of fruit fly locomotion

How does TREBLE perform when applied to biological data? First, we applied the TREBLE to fruit fly locomotor behavior collected from individual animals walking on an air-cushioned ball while exploring a virtual world (Haberkern et al. 2019). We extracted rotational, translational and slip velocity components from these data and applied the empirical window analysis outlined above. Structural variation analysis of behavior space revealed a minima at a window size of 180ms (Figure 2A; Figures S2A-C) while mean recurrence time was minimized at 140ms (Figure 2B; Figures S2D-F). Given that these metrics displayed only a small amount of variation between these two window sizes, we focused further analyses on a behavior space constructed using a window size of 160ms.

**Figure 2.**
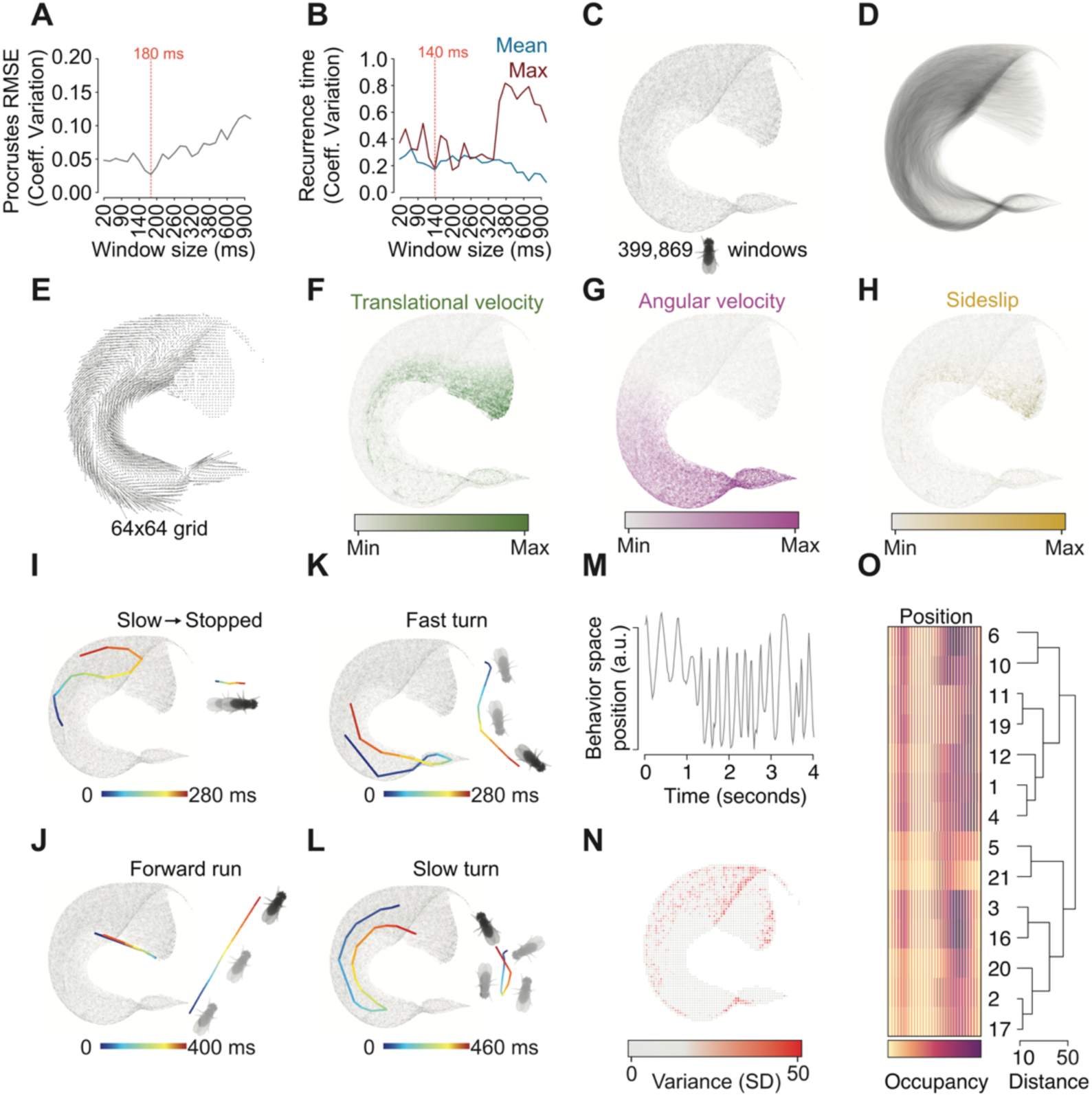
Analyzing fruit fly locomotion with TREBLE. **(A)** The coefficient of variation of Procrustes distance as a function of window size. The observed minimum at 180ms is denoted by the red dotted line. **(B)** Coefficient of variation for the mean (blue line) and maximum (i.e. highest proportion of time bins displaying recurrence; dark red line) recurrence times as a function of window size. The optimal tradeoff between the mean and maximum times is denoted at 140ms with a red dotted line. **(C)** Fly locomotor behavior space. Each point corresponds to a feature window (399,869 windows in total) as extracted from the 14 individual flies. **(D)** Pathways through the locomotor behavior space, produced by connecting temporally adjacent windows with partially transparent lines (as in Figure 1F). **(E)** Walking fruit fly behavior space represented as a vector field. Arrow direction and magnitude correspond to the angle and mean direction taken after visiting each bin. **(F)** The distribution of translational velocity across behavior space. Darker green corresponds to larger values of translational velocity. **(G)** The distribution of angular velocity across behavior space. Darker purple corresponds to larger values. **(H)** The distribution of side slip across behavior space. Darker gold corresponds to larger values. (**I)** Example bout of stopping. The pathway through behavior space is represented on the left. XY coordinates of the actual bout are plotted on the right. Both representations are colored by elapsed time. (**J-L)** Same as **(I)** but for fast turns, forward runs, and slow turns, respectively. **(M)** Sample trajectory through behavior space as represented by a 1-dimensional coordinate value. **(N)** Distribution of per-bin standard deviation across locomotor behavior space. Color corresponds to the variance of bin-wise visitation frequency across all 14 flies. **(O)** Hierarchical clustering of individual fly density maps. The dendrogram on the right represents the relationships between all 14 flies (numbered at the tips of each branch). Individual fly density maps are presented on the left in 1-dimension by converting the density map matrix into a single vector (presented in 2-dimensions in Figure S2J). Darker colors reflect more time spent (‘occupancy’) in a specific region.

We constructed a behavior space from the movements of 14 individual walking fruit flies (Figure 2C; 399,869 windows). Individual points in this space corresponded to unique windows with specific combinations of velocity features (Figure 2C). Connecting these points based on their temporal order revealed stereotyped paths in which individual flies repeated the same patterns of movement (Figure 2D; Figure S2G). As a result, the mean vector field produced by these was highly structured (Figure 2E). These pathways traversed regions of space defined by the input velocity features, meaning that position within the space could be used to infer the underlying pattern of movement (Figures 2F-H). Analyzing continuous movement through these regions demonstrated that individual behavioral bouts and sequences could be identified (Figures 2I-L; manually chosen). Finally, consistent with the notion that these sequences were recurrent (Figure 2B), plotting the positions of individual fly trajectories in behavior space over time reveals periodicity (Figure 2M, and data not shown). Taken together, these observations show how TREBLE can be used to identify repeated, intuitive, and interpretable patterns of behavior over time.

The capacity of TREBLE to co-embed many trials or individuals in the same space facilitates direct measurements of trial-to-trial or individual-to-individual variation. For example, we found that each of the 14 flies moved in grossly similar ways, as indicated by the fact that 83% of the behavior space was explored by all individuals, while less than 1% of the behavior spaces was explored by only one individual (Figures S2H-I). At the same time, individual trajectories varied greatly in how often they traversed different parts of the space. To measure this, we computed a probability density map for each individual (Figures S2J), and identified small, specific regions of behavior space that had the greatest variance between individuals (Figure 2N; Figures S2H-I). Hierarchical clustering of 1-d vectorized versions of these (Methods) maps revealed a variety of behavioral profiles across individuals (Figure 2O). Thus, in this case, behavior space was composed of movement types common to all individuals and could therefore act as template to compare the unique statistics of each.

### Detecting behavioral changes due to optogenetic manipulation

To test the ability of TREBLE to detect behavioral changes arising from an experimental manipulation, we analyzed locomotor behavior during optogenetic activation of gustatory sensory neurons expressing Gr5a, a sugar receptor (Haberkern et al. 2019) (Figure 3A; see Methods). To induce local search behavior, a 200 ms optogenetic stimulation was delivered every time a fly reached a pre-specified area and was repeated for every return visit. We processed nineteen optogenetic trials (1,110,025 windows) and co-embedded these windows with those from the control flies (described above) in the same behavior space. The movements contained within optogenetically stimulated trials overlapped extensively with those in control flies, with >97% of the space occupied by both datasets (Figures S2K-L). However, on average, the frequencies with which specific patterns emerged in the two datasets diverged dramatically (Figures 3B-C) and were unevenly distributed across behavior space (per bin Kruskal-Wallis test, see Methods; Figure 3D). These observations suggest that, while the structure of locomotion is conserved overall, these two groups display quite different temporal patterns of behavior.

**Figure 3.**
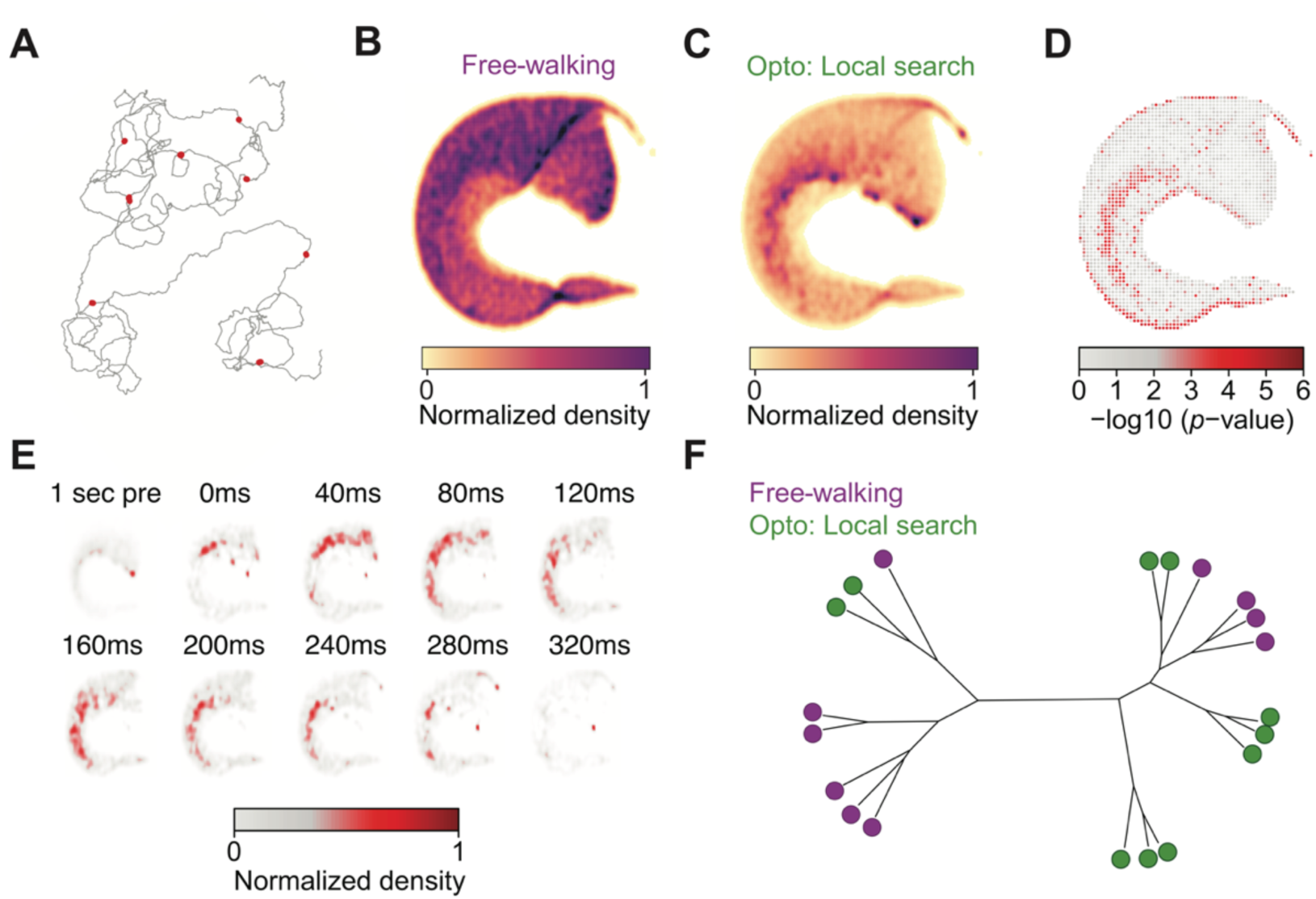
Identifying behavioral perturbations via optogenetic manipulation. **(A)** Example trajectory for a single optogenetic trial. The fly’s path through the virtual world is represented by the dark grey line. Red dots correspond to locations in which optogenetic stimulation was presented. **(B)** Average density map computed from the 14 free-walking control flies (darker color corresponds to more time spent in a specific region). **(C)** Average density map computed from the 19 optogenetically activated flies. **(D)** Bin-wise differences between control and optogenetically activated flies. Color corresponds to the significance (-log10 transformation of the *p-*value; Kruskal-Wallis test) of the differences in density between the two groups. Darker red corresponds to increasingly significant differences. **(E)** Time evolving responses to optogenetic stimulation in behavior space. Each space reflects a specific time window before (first space) and after stimulation (each subsequent space). Color corresponds to the average response to stimulation across all 19 flies (represented as normalized density; see methods). Windows span 1 second of time, beginning at the value represented above each space. **(F)** The behavioral relationships between all free-walking control and optogenetically activated flies. The tree represents the results of hierarchical clustering on the density maps of each individual fly (as in Figure 2O). Each branch tip is associated with an individual fly, group is represented by color (Controls = purple; Optogenetic trials = dark green).

To further explore this difference, we leveraged the continuous, time-resolved nature of TREBLE to map the evolution of average responses to neuronal activation over time (Figure 3E). Compared to pre-stimulus epochs, overall behavioral responses displayed an initial period of slowing (spanning from 0 to 120ms after stimulus offset), followed by increased turning (120ms to 280ms) (Figure 3E). These dynamics are consistent with known local search behaviors (Haberkern et al. 2019; Corfas et al. 2019). Despite these population-level patterns, we noted a surprising amount of behavioral heterogeneity amongst optogenetically activated flies, such that the behavior of some optogenetically activated flies overlapped with that of controls (Figure 3F; Figures S2J-M). Thus, TREBLE can assess the effects of a behavioral perturbation to detect both population and individual level variation.

### Identifying structure in more complex feature sets: Locomotion of *Drosophila* larvae

Thus far we have used to TREBLE to analyze behaviors as changes in centroid velocity components. However, pose estimation methods have made analyses of other behavioral features, such as posture or limb movement, increasingly common (Pereira et al. 2020; Mathis et al. 2020). As such, pose estimation methods typically represent behavior in multi-dimensional spaces that may or may not include explicit velocities. With this in mind, we next assessed the ability of the TREBLE framework to capture a high-dimensional combination of postural and velocity features describing larval *Drosophila* locomotion.

Locomotion in *Drosophila* larvae is characterized by stereotyped changes in size and posture correlated with peristaltic movements and bending that produce forward and backward translation and turning (Clark et al. 2015). To capture this, we tracked *Drosophila* larvae via machine vision and at each time point calculated 11 complex postural and velocity features following established methods (Figure 4A; n = 72) (Risse et al. 2014, 2017). Given that certain of these features may be correlated, we performed principal component analysis and found that 8 components were sufficient to explain over 90% of the observed variance (Figure S3D). These 8 PCs were then used to run the iterative window procedure, resulting in an optimal window size of 800ms (Figures S3A-C). This produced a 2D behavior space that revealed a strongly oscillatory region (Figure 4B) with highly directional movement (Figure 4C). Analyzing the distribution of the input features revealed that this oscillator corresponded strongly with features of peristaltic locomotion (Figures 4D-G). Specifically, the oscillator appeared to switch between a regime of lower velocity paired with an increase in the area of the animal’s shape (i.e. scrunching) and higher velocity with lower overall area (stretching) (Figures 4D-G). When oscillating, these regimes produced an individual ‘run’ of crawling (Clark et al. 2018).

**Figure 4.**
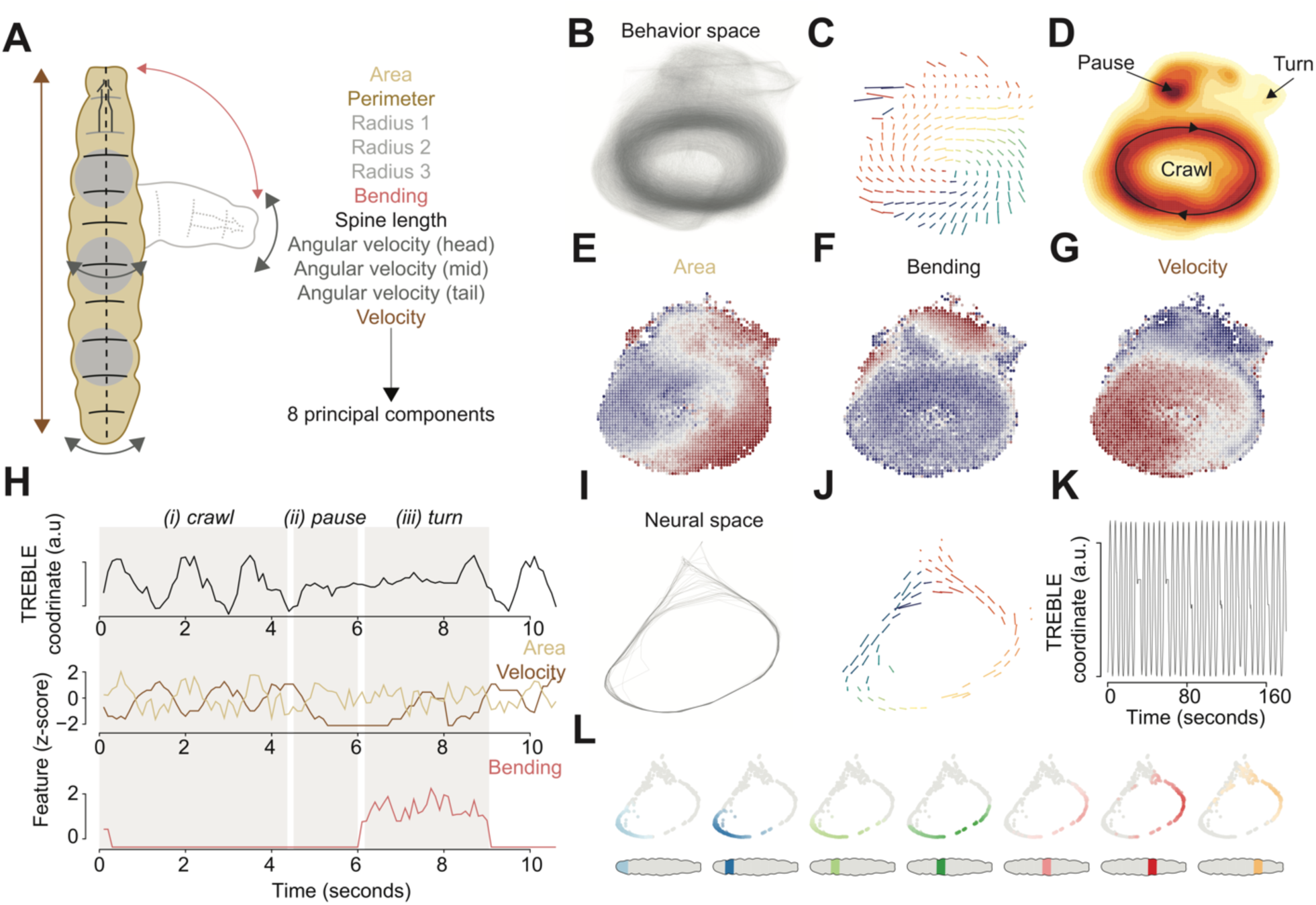
Larval *Drosophila* crawling dynamics. **(A)** Cartoon of larval *Drosophila* movement and accompanying features used for analysis. **(B)** Pathways through the larval locomotor behavior space (n = 72 larvae). **(C)** The mean vector field of larval locomotor space. Direction and size of arrow correspond to the mean movement through a given bin in space. Color denotes the angle of the arrow. **(D)** Probability density function of larval locomotor space plotted as a heatmap. Behaviors annotated qualitatively. **(E-G)** The distribution of area **(E)**, bending **(F)**, and velocity **(G)** as a function of larval behavior space (z-scores). Blue corresponds to negative values, red represents positive values. **(H)** Example distribution of behavior space position (TREBLE coordinate; a.u.), area (z-score; yellow), velocity (z-score; brown), and bending (z-score; read) over a ∼10 second epoch of behavior. The approximate durations of behavioral states are annotated via grey shading and labeled above (i: crawl, ii: pause, iii: turn). **(I)** Pathways through larval neural space (plotted here is an example distribution of a single trial; full space in Figure S3I). **(J)** Larval neural space plotted as a mean vector field. **(K)** Example of oscillating position in neural space over a 160 second period of time. **(L)** Probability density functions of motor neuron activity across behavior space. The plots proceed from posterior (leftmost) to anterior (rightmost) as denoted by the cartoon larvae plotted below (with approximate location of motor neuron segment colored).

The upper portion of the space, outside the oscillating domain, was split between a region of increased bending and another associated with pausing (Figure 4D). Comparing the distribution of features over time to movement in the TREBLE space highlighted the structured relationships between these features (Figure 4H). During crawling (denoted by ‘i’ in Figure 4H) velocity and area oscillate inversely. When a turn is initiated, the animal first pauses (ii), decreasing velocity and stabilizing area, and then begins bending with a corresponding shift in position in behavior space (iii). After the turn is complete, oscillatory crawling begins again (Figure 4H). Overall, oscillatory waves occur with a period of around 1 second (Figure S3E), matching previous observations of *Drosophila* larval crawling (Heckscher et al. 2012). TREBLE therefore captured the common structural and temporal elements of the *Drosophila* larval ethogram (Clark et al. 2016) within a single behavior space.

The recurrent properties of crawling mirror waves of motor neuron activity (Tastekin et al. 2018; Clark et al. 2018). We therefore wondered if using TREBLE to analyze the output of motor neurons might also yield a corresponding oscillator describing neural dynamics. We used calcium imaging data from fictively crawling larvae (Tastekin et al. 2018) to construct a ‘neural space’ from the activity of seven motor neuron classes (Figure 4I; Figures S3F-K).

As with the behavior space, we found that motor neuron activity yielded an oscillator (Figure 4I) with directional (Figure 4J) and stereotyped movement (Figure 4K). Bouts of crawling are associated with waves of neural activity, originating in posterior neurons and propagating in the anterior direction (Tastekin et al. 2018; Clark et al. 2018). Plotting the peak activity of each motor neuron type in the neural space recapitulated this observation (Figure 4L). Peak activity of the posterior neurons occurred in the lower left-hand portion of the space, followed by spatially sequential peaks of the more anterior motor neurons (Figure 4L) moving in the same direction described the mean vector field of the space (Figure 4J). These observations reveal that both the behavioral output and neural dynamics underlying *Drosophila* larval locomotion can be captured in a common oscillatory framework. In addition, these analyses show that TREBLE can applied to find structure in higher-dimensional neural and behavior data.

### Identifying structure in more complex feature sets: Mouse pose dynamics

We next sought to generalize TREBLE to a much more complex form of behavioral data, mouse movements in three dimensions (from Markowitz et al. 2018). To do so, we analyzed mouse behavior measured with 3D imaging via the MoSeq pipeline (Wiltschko et al. 2015; Markowitz et al. 2018). In this pipeline, freely moving mice were imaged in an arena using three orthogonal cameras at 30Hz (Wiltschko et al. 2015; Markowitz et al. 2018). These video streams were then processed to produce 17 behavioral features (Figure 5A) representing mouse movement in three dimensions (Wiltschko et al. 2015; Markowitz et al. 2018) which we then used as input to the iterative window size procedure. A window size of 130ms was chosen (Figure S4A-C), yielding a behavior space that organized aspects of posture and movement into a recognizable, and recurrent, structure (Figures 5B-J). Analyzing the distribution of input features in behavior space (Figures 5E-J) allowed us to identify portions of the space related to characteristic behaviors such as walking, scrunching, rearing, and pausing (Figures 5C-D). These designations were corroborated by calculating the average 3D pose of animals as a function of position in behavior space (Figure S4D). These observations indicate that TREBLE can be used to identify structure in high-dimensional pose estimation datasets.

**Figure 5.**
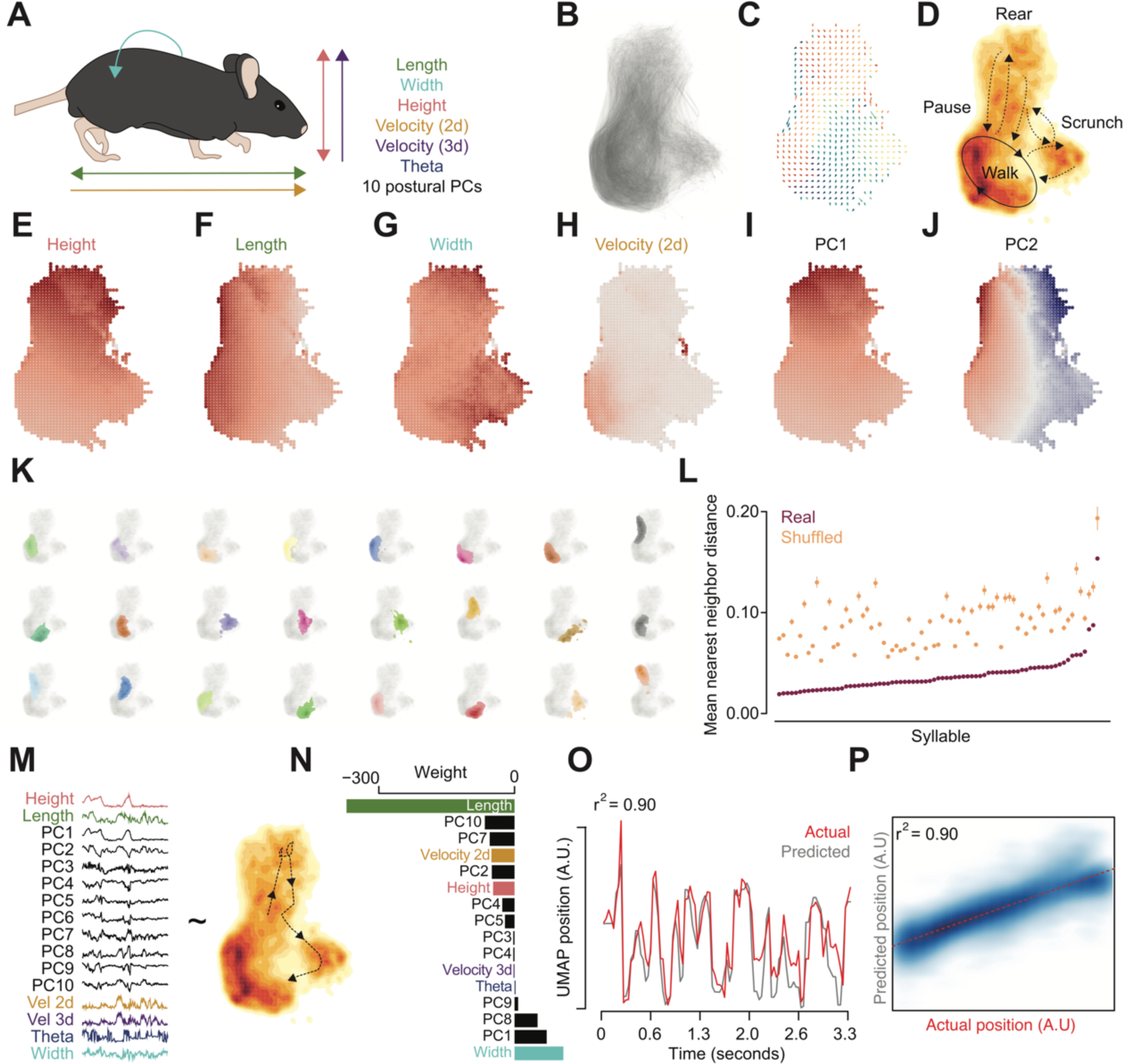
Analyzing 3D pose dynamics in mice. **(A)** Cartoon of mouse 3D movement and accompanying features used for analysis. **(B)** Pathways through mouse 3D pose space (n = 8 mice). **(C)** The mean vector field of mouse 3D pose space. Direction and size of arrow correspond to the mean movement through a given bin in space. Color denotes the angle of the arrow. **(D)** Probability density function of mouse 3D pose space plotted as a heatmap. Behaviors annotated qualitatively. **(E-J)** The distribution of height **(E)**, length **(F)**, width **(G)**, 2D velocity **(H)**, postural PC1 **(I)**, and postural PC2 **(J)** as a function of larval behavior space. **(E-H)** Color ranges from grey (minimum value) to red (maximum). **(I-J)** Blue corresponds to negative values, red represents positive values. **(K)** The distributions of the 24 most common behavioral syllables (as identified by MoSeq) in behavior space. A probability density function across behavior space was computed for each syllable and then plotted in color on top of the full behavior space (in grey; see Methods). **(L)** The distribution of mean nearest-neighbors distance between points in behavior space for all syllables (n = 43). Purple denotes the observed values. Orange corresponds to the mean and distribution (lines; lower (25^th^ percentile) and upper (75^th^ percentile) hinges of a boxplot) of shuffled data (10,000 permutations). **(M)** Visualization of the variables used to construct the regularized generalized linear model. Features are represented by example time series (left) and were compared to movement through behavior space (right). **(N)** Barplot of coefficient weights from the final model, sorted by weight and colored to match the example time series in **(M)**. **(O)** Comparison of the actual position in behavior space (grey) to the prediction from the final GLM (red) for an example ∼3 second time period. **(P)** Smoothed scatterplot comparing observed and predicted behavior space positions for the full dataset. Darker blue denotes greater density of points. Dashed red line corresponds to the fit of a regression between observed and predicted values.

In addition to measurements of 3D movement, the MoSeq pipeline also identifies how discrete elements of mouse behavior - behavioral ‘syllables’ – are sequenced over time (Wiltschko et al. 2015; Markowitz et al. 2018). What is the relationship between the continuous representation of 3D behavior provided by TREBLE and the discrete output of the MoSeq pipeline (Markowitz et al. 2018)? Annotating the TREBLE behavior space with the individual syllables identified by MoSeq revealed that the distributions of individual syllables appeared highly compact (Figure 5K; Figure S4E). Syllables produced by MoSeq corresponded to distinct and identifiable behavioral states identified by TREBLE, such as walking (Figure 5K; yellow and blue syllables; first row, fourth and fifth columns) and scrunching (Figure 5K; purple syllable; second row, third column). Furthermore, these distributions were non-random. By comparing the mean nearest neighbor distance for each syllable to a shuffled distribution, we found that the mean distance between points was significantly lower than expected by chance (*p* < 0.0001; permutation test) for all syllables (see Methods). Therefore, TREBLE and MoSeq capture complementary aspects of behavior - both discrete and continuous – albeit using very broadly different statistical frameworks.

Finally, we wondered whether TREBLE was truly capturing the majority of behavioral variation present in this complex dataset. To address this, we created a regularized generalized linear model (GLM) comparing the relationship of the input features to the position in TREBLE behavior space over time (Figure 5M; see Methods). We found that the GLM captured over 90% of the variance in the data, and pose features such as length and width, and related postural principal components, contributed strongly to the model (Figures 5N-P). Position in the TREBLE space could be predicted with a substantial degree of accuracy from the feature set (Figures 5O-P), predictability that was consistent across individual trials (Figure S4F). We therefore conclude that TREBLE is able to explain a substantial portion of behavioral variation even in complex feature sets and can robustly represent the 3D dynamics of animal movement in a low-dimensional, continuous framework.

### Using TREBLE to characterize neural encoding of behavior

A common, and often difficult, goal in neuroscience is to relate neural activity to behavior. We therefore used TREBLE to identify behavioral coding in 794 medial entorhinal cortical (MEC) neurons recorded during the foraging trials presented above (n = 14 mice, 327 trials) (Hardcastle et al. 2017) (Figure 6A). The MEC is hypothesized to support navigation and contains a population of functionally defined neurons that encode a variety of behavioral variables such as an animal’s position in external space (e.g. grid and border cells), head direction, and running speed (Kropff et al. 2015; Sargolini et al. 2006; Solstad et al. 2008; Hafting et al. 2005).

**Figure 6.**
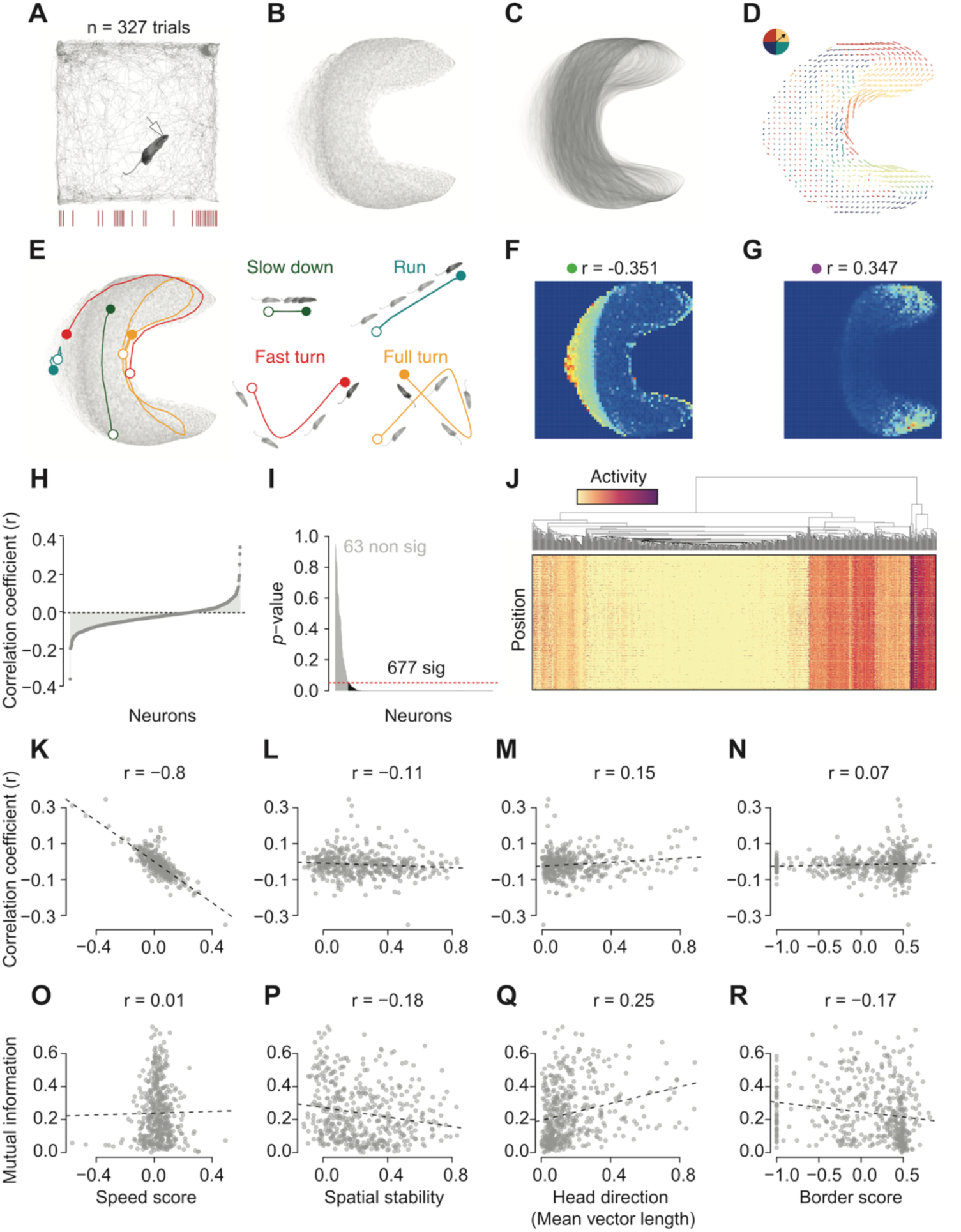
Mapping mouse navigational coding. **(A)** Example trajectory for of a foraging mouse with paired spike recordings via an implanted electrode (drawn). **(B)** Mouse locomotor behavior space, each point corresponds to a temporal window. **(C)** Pathways through mouse locomotor behavior space, produced by connecting temporally adjacent windows with partially transparent lines. **(D)** Mouse locomotor behavior space represented as a vector field. Arrow direction and magnitude correspond to the angle and mean direction taken after visiting each bin. Arrows are colored by the degree of the direction vector (corresponding to circle in upper left-hand corner). **(E)** Example behavior bouts decoded from behavior space. Pathways through behavior space are plotted on the left. Bout starts are indicated by the open circle, ends by the closed circle. Positions in real XY space are plotted on the right. **(F)** Example 2-d tuning curve of a neuron that is negatively correlated with behavior space (activity measured as spikes/second; labeled with a green dot for reference in later figures). Colors range from blue to red, corresponding to minimum activity (blue) to maximum activity (red). Pearson correlation (r) is denoted above the plot. **(G)** Example 2-d tuning curve of a neuron that is positively correlated with behavior space (activity measured as spikes/second; labeled with a purple dot for reference in later figures). Colors range from blue to red, corresponding to minimum activity (blue) to maximum activity (red). Pearson correlation (r) is denoted above the plot. **(H)** Distribution of correlation coefficients between neural activity and behavior space position for all MEC neurons (activity measured as spikes/second). **(I)** Distribution of *p-*values (Bonferroni corrected) resulting from permutation tests (10,000 shuffles) of the correlations plotted in **(h)**. The red dotted line indicates *p =* 0.05. **(J)** Clustered 2-d tuning curves for all neurons. The heatmap corresponds to a linearized version of the 2-d tuning curve, colored by average activity as a function of position in behavior space. **(K)** Scatterplot of the relationship between the correlation coefficient of the relationship between behavior space and neural activity (y-axis) and speed score (x-axis). Pearson correlation (r) is denoted above the plot. **(L)** Scatterplot of the relationship between the correlation coefficient and spatial stability. **(M)** Scatterplot of the relationship between the correlation coefficient and head direction (HD) mean vector length. **(N)** Scatterplot of the relationship between the correlation coefficient and border score. **(O)** Scatterplot of the relationship between the mutual information (MI) of behavior space and neural activity (y-axis) and speed score (x-axis). **(P)** Scatterplot of the relationship between MI and spatial stability. **(Q)** Scatterplot of the relationship between MI and head direction (HD) mean vector length. **(R)** Scatterplot of the relationship between MI and border score.

We extracted time-varying velocity measures from positional data of individual mice as they foraged for randomly scattered food rewards in a 1m by 1m box (Figure 6A). We then collected windows from these data using the iterative selection procedure to choose a window size of 400ms (Figure S5A-F). As we previously observed with the correlated random walk and *Drosophila* spaces, the resulting behavior space ordered points by velocity features (Figures S5G-L) and connected them with continuous and directionally recurrent pathways (Figures 6B-D). Moreover, distinct behavioral bouts – such as running, turning, and stopping – could be easily decoded from the space and mapped back onto real XY coordinates (Figure 6E).

We related MEC neuron activity to movement in the behavior space defined by TREBLE. To do so, we first created a 2-dimensional behavioral tuning curve for each neuron. Each cell’s average activity was mapped as a function of the animal’s position in behavior space, revealing a wide variety of patterns (Figure S5M). Correlation coefficients were then calculated by comparing each neuron’s activity with a 1-dimensional representation of behavior space position (64 x 64 grid: Methods). Strongly positive or negative correlations thus reflect that a given’s neuron activity is increased in different portions of behavior space (Figure 6F-H), of which a number were identified. Consistent with the MEC’s role in navigation, permutation tests revealed that these correlations were overwhelmingly non-random, with only 63 of the 794 MEC neurons displaying non-significant relationships with locomotor behavior after multiple test correction (Figure 6I). Furthermore, clustering neural activity patterns across behavior space revealed multiple distinct types of relationships between MEC neurons and locomotor behavior (Figure 6J; Figure S5M). Most neurons were active in relatively small and specific regions of behavior space, suggesting that they encode information relevant to particular locomotor movements (Figure S5M). Other neurons displayed distributed activity across behavior space, consistent with less selective coding of locomotor information (Figure S5M). Thus, these various patterns can be considered 2-d tuning curves, representing the relationship between neural activity and behavior for each neuron.

To what extent do these patterns reflect previously described representations of navigational coding in the MEC? To address this, we intersected TREBLE correlations with other commonly calculated MEC coding variables (speed, head direction, spatial stability, border proximity; see Methods). Unsurprisingly, we found that behavior space correlated with speed coding (Figure 6K) compared to the other measures (Figures 6L-N). However, a more precise measure of coding capacity requires a comparison between the amount of information jointly shared between position in behavior space, speed score, and neural activity. To do this, we computed the mutual information (MI) between TREBLE position and neural activity for each neuron and then related these measures to the commonly calculated MEC coding variable (Figures 6O-R). Higher MI values here indicate that spatial coordinates in behavior space contain more information about a given neuron’s activity, reflecting stronger behavioral coding. Strikingly, MI was uncorrelated with speed score (Figure 6O), indicating that encoding of locomotor behavior was not solely related to variation in speed. This raised the possibility that these neurons may encode information about other locomotor variables including longer timescale and more complex maneuvers.

### Using TREBLE to identify temporal variation in the neural coding of behavior

Neural activity can relate to continuous behaviors on a variety of timescales, the discovery of which is an area of active interest (Datta et al. 2019; Krakauer et al. 2017; Glaser & Kording 2016). We therefore wanted to test the ability of TREBLE to provide a path for relating neural activity with behavior and took advantage of the apparent rich capacity of MEC neurons to encode locomotor behavior over a variety of timescales.

To do this, we first computed the cross-correlation of TREBLE position and activity for each neuron (Figures 7A-C). This analysis identifies when a neuron’s activity correlates with position in behavior space by calculating correlations between the two variables using a sliding window centered around a temporal offset of zero (i.e. simultaneity). We found a variety of temporal offsets in the cross-correlation distributions for each neuron. Peak correlations could both precede (Figure 7A) and follow (Figure 7B) changes in behavior. Clustering the distribution of these cross-correlation coefficients revealed a variety of neural classes, including negative, positive, and even mixed negative and positive cross-correlation profiles (Figure 7C; Figure S6A). Notably, cross-correlation peaks could be of varying widths; some were tightly centered (e.g. Figures 7A-B) while others displayed correlations lasting seconds (Figure S6A). Therefore, in addition to their instantaneous relationships, the activity of MEC neurons can covary with behavior over a range of temporal scales and offsets.

**Figure 7.**
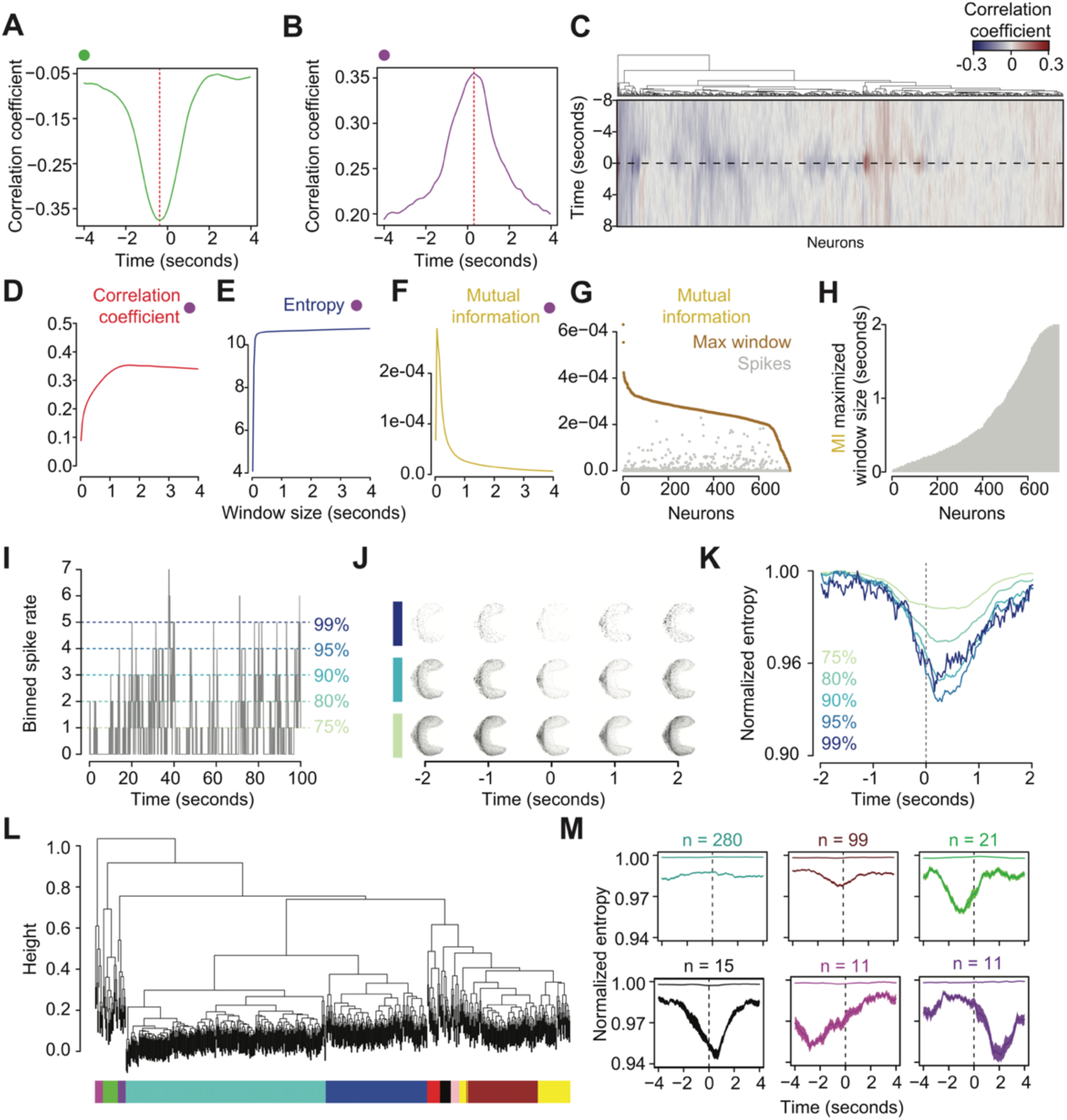
Temporal variation in mouse navigational coding. **(A)** Cross-correlation distribution of a neuron that is negatively correlated with behavior space (same as in **6F**). **(B)** Cross-correlation distribution of a neuron that is positively correlated with behavior space (same as in **6G**). **(C)** Hierarchical clustering of cross-correlation distributions across all neurons. Color corresponds to the correlation coefficient, blue indicating negative and red indicating positive correlations. The y-axis represents time before and after instantaneity, ranging from 8 seconds before to 8 seconds after. The x-axis corresponds to the relationships of all MEC cells based on their cross-correlations with behavior (which are shown via the colored heatmap). In essence, each column contains the same information as **7A-B** but here represents the distributions using variation in color over time, rather than a 1-d tuning curve. **(D)** Correlation coefficient distribution as a function of varied window sizes for the neuron plotted in **6G** and **7B**. **(E-F)** same as **(D)** but for entropy **(E)** and mutual information (MI). **(G)** Distribution of MI for all neurons, comparing the maximally informative window size (in gold) and that measured from spikes (grey). **(H)** Barplot of the distribution of MI maximized window sizes for all neurons. **(I)** Example neural activity trace (using MI maximized window size) and corresponding percentile cutoffs (at 75%, 80%, 90%, 95%, and 99%). **(J)** Temporal distributions in behavior space (1 second windows) surrounding peaks in activity (75%, 90%, and 99% cutoffs). **(K)** Entropy distributions of surrounding activity bouts corresponding to each percentile (labeled and colored on plot). Normalized entropy measures are shown to be able to compare across percentiles with different sample sizes. **(L)** Hierarchical clustering of entropy distributions across all neurons. Colored bars indicate clusters as measured using dynamic tree trimming (see Methods). **(M)** Example mean entropy distributions for four clusters identified using dynamic tree trimming. Plotted are the distributions for the 75% (top) and 99% (bottom) cutoffs. Mean and standard error are plotted as well as sample sizes (above).

Given this observation we hypothesized that individual MEC neurons may most reliably encode behavior over specific timescales. To test this, we calculated each neuron’s activity over a range of temporal windows and measured their associations with behavior space via correlation, mutual information (MI), and entropy (i.e. how narrowly neural activity was distributed across behavior space) (Figures 7D-F). For most neurons, the strength of the correlation between activity and behavior as well as entropy plateaued (Figures 7D, 7F) while MI displayed a clear, and at times very specific, maximum value as a function of temporal window size (Figure 7F; Figure S6B). This pattern suggested that, given temporal binning that maximized MI, neural activity could be more specifically associated with distinct regions in behavior space. This was a general pattern: across all neurons the MI maximized window size encoded more information than instantaneous spikes (Figure 7F). The distribution of these MI maximized window sizes varied broadly, ranging from instantaneous spiking to window sizes up to 2 seconds, reflecting the diversity of patterns observed in the cross-correlation analyses (Figure 7G; Figure 7C). MEC neurons thus possess specific and, in some cases, finely tuned temporal scales over which their activity most informatively encodes movement.

Finally, we asked whether these temporal relationships might be further refined by considering how the magnitude of each neuron’s activity constrains its behavioral coding. Do extrema in neuronal firing rates reflect specific behaviors? To answer this question, we calculated percentile cutoffs for each neuron’s activity across the trial using the MI maximized window sizes (Figure 7I). We then examined whether increasing activity rates were associated with consistent changes in movement through behavior space, indicating increasing behavioral specificity. To do so, we analyzed the distribution of movements within behavior space by computing the entropy of the distribution associated with neural activity bouts above each cutoff (Figures 7J-K). In this metric, lower entropy values correspond to increased stereotypy in behavior space. For many neurons, as activity increased the corresponding behavior space entropy decreased, meaning that high levels of neuronal activity tended to be associated with specific behavioral patterns (Figure 7K; Figure S6C). Hierarchical clustering of these entropy distributions separated neurons into coherent groups with temporally similar profiles (Figures 7L-M). Notably, a number of clusters displayed entropy minima that were significantly offset from zero, meaning that peaks in neural activity could either precede or follow changes in behavior (Figure 7M; Figure S6C). Taken together, these results demonstrate that MEC neurons can encode locomotor information on temporally varying scales, both before and after changes in behaviors occur. Moreover, these analyses demonstrate how TREBLE can be used to uncover the relationship between neuron activity and continuous behavior across a rich neural dataset from freely moving animals.

## Discussion

Technological developments have led to substantial shifts in how researchers are attempting to unlock the development, expression, and evolution of animal behavior. Facilitated by high throughput data analysis, a number of ambitious conceptual projects have recently been proposed. These include considering behavior as a natural extension of physics (Brown & de Bivort 2018), attempting to capture many species’ complete behavioral repertoires (Anderson & Perona 2014), or even reconciling the two dominant lineages of behavioral research, comparative psychology and ethology (Datta et al. 2019). Achieving these goals requires the study of individuals across a diversity of animal species, in both controlled and naturalistic contexts, and in conjunction with other data streams, critically including neural activity. Given this, behavioral analytical methods that are statistically robust, broadly applicable, and easily linked to the function of nervous systems are of general interest. Here we show how TREBLE can produce low-dimensional representations of behavior that capture key features of the temporal structure of animal movements and poses, allow one to visualize the phenotypic consequences of perturbations, and reveal relationships to neural activity. By analogy with genomics, these “heatmaps” can constitute analytical endpoints, but can also guide the subsequent development of more targeted hypotheses.

All analytical methods for parsing behavior inherently make tradeoffs between computational ease, temporal resolution, and generalizability. TREBLE does not require extensive model fitting, is agnostic to feature type, and can accommodate many millions of data points. Moreover, we describe how the key parameters needed, such as window size, can be systematically explored and rationally chosen to capture behavioral structure of interest. By cutting behavior into defined temporal windows that are extremely densely sampled, TREBLE simultaneously captures both long and short temporal structures in behavior. Furthermore, representation of these can be handled flexibly. For example, here we represent temporal structure in two ways: 1) as continuously varying time series and 2) as information theoretic summary measures of entire distributions over time (using entropy and mutual information). It is also possible for users to discretize TREBLE’s output simply by segmenting movement through behavior space into sub regions. Finally, TREBLE is generalizable in three different ways. First, as we demonstrate, the framework can be applied to a variety of movement sources, including synthetic trajectories and multiple animal species. Second, the position of TREBLE in analysis pipelines is adjustable. While TREBLE can be implemented as a standalone method for behavioral analysis from beginning to end, it can also be used as a screening tool for identifying statistically significant behavioral variation to be followed by other approaches specifically tailored to the question of interest. Finally, as we show using optogenetic activation in flies and neural recordings in mouse MEC, TREBLE can be used to find meaningful links between nervous systems and behavior in a variety of contexts. This versatile nature of the TREBLE framework therefore opens up the possibility for customizable high-throughput behavioral analyses.

TREBLE can uncover otherwise opaque behavioral variation by directly comparing behavioral data across many individuals and conditions. Here, we show that free walking fruit flies possess substantial variation in locomotor dynamics and that this variation is associated with specific regions of behavior space. Surprisingly, we find that this variation in free walking behavior overlaps with the diversity of responses observed after an optogenetic perturbation. A limited set of pre-determined behavioral measurements, as would have been used typically, may have missed this variation. Similarly, even unbiased statistical methods, without clear grounds for comparison across animals or experimental conditions, would have been unlikely to have detected these patterns. This highlights the benefit of using unsupervised statistical descriptions such as TREBLE for assessing the structure of behavior across diverse individuals. Furthermore, we envision that TREBLE might be employed on control data sets to measure variability across baseline individuals in order to facilitate statistical power calculations across the full range of behavioral dynamics that might be present. Such catalogs of variability could be thus be developed across species’ wildtype behavioral repertoires, allowing researchers to better account for inborn variance or threshold effects in applications such as genetic mapping, behavioral manipulations, and neural perturbations (Lopez-Alonso et al. 2015; Ayroles et al. 2015; Buchanan et al. 2015).

Associating neural activity with continuous behavior presents two key challenges. First, identifying the specific aspects of behavior that activity corresponds to is difficult to determine in the context of a large behavior repertoire. Second, the temporal scales over which this relationship occurs can be hard to know a priori. Here, we show how TREBLE can be used to address these issues by characterizing the oscillatory patterns of motor neuron activity during larval *Drosophila* crawling and the coding of locomotor behavior in mouse medial entorhinal cortex (MEC) neurons as proofs of concept.

In the latter case, we find that TREBLE captures known components of speed coding in MEC neurons while also uncovering extensive variation in temporal and rate-based coding. Previous work has found that MEC neurons prospectively code for changes in an animal’s speed and position ∼50-80ms in the future (Kropff et al. 2015). Our analyses add to these findings, suggesting that MEC neurons can encode changes both before *and* after behavioral events, doing so across a variety of timescales. These patterns may reflect the presence a multitude of temporal encoding strategies for behavior in the MEC, similar to what has been seen in basal ganglia movement selection (Markowitz et al. 2018; Jin & Costa 2015), orofacial rhythms (Moore et al. 2013), and whole-brain activity during *C. elegans* locomotion (Kaplan et al. 2020). We speculate that this may arise from some MEC neurons encoding short time-scale behavioral events (e.g. turning left or right) while others update based on longer-term navigational behaviors such as goal-oriented searching or foraging. These results demonstrate how TREBLE can be leveraged to identify specific aspects of behavior that are associated with the activity of individual neurons and to uncover the temporal structure of such relationships. More broadly, we anticipate that TREBLE may be useful in uncovering the statistical structure of behaviorally relevant activity across many diverse populations of neurons.

There are a number of areas in which TREBLE may be further developed and employed. First, movement through behavior space may be classified on timescale that are longer than those presented here, affording descriptions of behavioral state over time or the emergence of behavior in development. Second, the ability of TREBLE to co-embed multiple individuals with temporal resolution may make it particularly amenable to studying social and collective behavioral dynamics. Third, by readily capturing variability in behavior, TREBLE may be amenable to exploring differences across individuals arising from factors imbuing variance such as reaction norms, behavioral syndromes, and environmental or genetic variation. Finally, it is especially intriguing to consider how TREBLE may be further leveraged to jointly infer neural coding principles by applying the framework to the structure of behavioral and neural dynamics in parallel.

## Acknowledgements

We thank members of the Clandinin lab and Kiah Hardcastle for useful discussion and input. We are grateful to Jeffrey Markowitz and Sandeep Robert Datta for providing the mouse 3D pose dataset. This work was supported by the NIH (U19NS104655; T.R.C.) (F31; A.C.R.), the Simons Foundation (T.R.C., L.M.G.), and the Stanford School of Medicine Deans Office (R.A.Y.).

## Author contributions

R.A.Y. and T.R.C. conceived the study. R.A.Y. wrote the TREBLE code and performed data analysis with input from L.M.G on mouse behavior and MEC neuronal recordings. A.C.R collected the larval *Drosophila* crawling data. R.A.Y., L.M.G., and T.R.C. prepared the manuscript.

## Methods

### Datasets

Correlated random walks were produced using the `TrajGenerate` function in the `trajr R` package (McLean 2018). Ten replicate walks were produced per parameter tested (e.g. window size), sampled at a rate of 100 frames per second and consisting of 10,000 frames. All walks were generated using the same underlying angular and linear error distributions (Normal distribution; Mean = trajectory length; Standard deviation = 0.5 (angular)/0.2 (linear)).

Details of the *Drosophila* walking dataset can be found in Haberkern et al. 2019. Briefly, animals in the free-walking dataset (‘WTB hybrid’ genotype) were allowed to explore a circular matt acrylic platform (radius 11.4 cm) surrounded by a siliconized acrylic cylinder to prevent climbing. Videos were recorded from 120 cm above the platform using a Flea3 camera at 12.3 Hz with a spatial resolution ∼40 pixel/cm. Trials lasted 10 minutes after a 1-2 minute acclimation period.

The local search data were produced from female flies possessing optogenetically-accessible sugar receptor neurons (Gr64f-Gal4 > ChrimsonR). Animals were allowed to explore a virtual landscape consisting of distributed ‘cone forest’ while walking on a circular treadmill. Treadmill movement data (fly’s position and velocity) were collected at 360 Hz. Optogenetic stimulation was triggered whenever a fly crossed within a 10mm radius of a given cone and persisted for 200ms.

For the *Drosophila* larvae analyses, embryos were collected for 1 h on standard 3.0% agar molasses collection caps covered with a thin layer of wet yeast. Twenty-four hours later, hatched embryos were transferred to standard cornmeal fly food. After forty-eight hours (L2 larval stage), animals were collected and transferred to a Petri dish with1.2% agar and relocated to a behavioral room kept at 23°C and 60% humidity. Ten to fifteen minutes after acclimation to the room, groups of 5 to 10 larvae were transferred to a 30 x 30 cm 1.2% agar arena. After 15 to 30 seconds, locomotion was recorded using a FIM imaging system (Risse et al 2013, https://www.unimuenster.de) at 10 fps for 5 minutes. The FIM system was equipped with an azA2040-25gm (Basler) camera and a LM16HC-SW (Kowa) lens. Individual larvae were then tracked using FIMtrack software (Risse et al 2013).

Details of the mouse 3D pose dataset can be found in Marowitz et al. 2018 and Wiltschko et al. 2015. Depth and position were collected using a Microsoft Kinect V2 while individual mice explored a circular arena (collected at 30Hz; 512×424 pixels frame size). The mouse’s center and orientation were estimated using an ellipse fit. An 80 x 80 pixel box was then drawn around the mouse and used to rotate the frame so that the mouse was always facing the righthand side. These cropped and adjusted frames were then used as input for principal component analysis from which the top 10 of these postural PCs were used for downstream analyses.

Details of the mouse behavior and MEC dataset can be found in Hardcastle et al. 2017. Behavior and neural recordings were collected from two cohorts of adult wildtype mice: 5 male and 2 female C57BL/6 mice (405 MEC cells recorded) and 7 male C57BL/6J:129SVEV mice (389 MEC cells). Two polymide-coated platinum iridium 90%-10% tetrodes were implanted in each mouse prior to the experimental period. Behavioral data were collected approximately one week after surgery in large open environments with black walls containing chocolate flavored cereal to induce foraging (sizes varying; see Hardcastle et al. 2017 for details). The majority of recording sessions lasted between 30-35 minutes with a small number ranging in time between 12 and 122 minutes. During each session, position, head direction, and running speed were recorded every 20ms and single unite spikes were recorded at a 10kHz sampling rate.

### Generation of correlated random walk behavior spaces

The trajectories and velocity distributions of replicate correlated random walks were extracted using the custom function `iterative_umap`. First, a novel walk trajectory was generated using `TrajGenerate` (as described above). Instantaneous velocity components were then calculated (translation, angular velocity, sideways velocity) and sampled using windows of a specified width (denoted here as `w`) and step size (denoted here as `s`). The sampling procedure was as follows.

First, given frame `i`, the velocity components were extracted for frames `i:i+w`. Angular and sideways velocity values were normalized to the first frame in the window so that each velocity vector to originate from zero. Since we weren’t concerned with information pertaining to turn direction (i.e. left vs. right) the proceeding velocity values were then adjusted so that the second frame was always positive. The resulting velocity vectors were linearized and concatenated, resulting in a single vector of length `3w`. This procedure was then repeated iteratively every `s` frames for the length of the trial (`t`). After each iteration the concatenated vectors were appended to a matrix with `3w` rows to ultimately create `t/s` columns.

This procedure yielded a library of densely sampled velocity fragments corresponding to behavioral dynamics for each moment in the trial. The `R` implementation of the `UMAP` algorithm (McInnes et al. 2018) was then used to embed these fragments into a low-dimensional behavior space (each point in space corresponding to a window). The resulting space thus provided a 2-dimensional xy position for each point/window in the trajectory. For downstream analyses we computed simplified positional information for each space using the `bin_space` function. This function decomposes behavior space into a grid of desired size (*n* bins x *n* bins). For example, for a desired grid of 16 bins x 16 bins, a new set of xy coordinates will be calculated corresponding to 16 intervals spanning the minimum/maximum xy coordinates in the original space. The position of each point in the space will then be compared to the new coordinates and associated with the closest bin, in this example case generating a reduced set of 16×16, or 256, unique bins.

### Analyzing correlated random walk parameter space

We assessed the relationship of parameter choice and behavior space structure by sweeping the two main window sampling parameters: window width and step size. We examined 24 different window widths, ranging from 10ms to 2 seconds (10-40ms sampled at 10ms steps; 50ms-2 seconds sampled at 100ms steps). Ten correlated random walk replicates were generated for each window size and then processed and embedded into behavior space using the `iterative_umap` function. We also explored the effect of step size by producing replicate behavior spaces from trajectories sampled at step sizes varying from 10 ms (i.e. 1 frame) to 1 second, separated by 100 millisecond intervals. Again, for each step size 10 correlated random walk replicates were generated, processed, and embedded into behavior space using the `iterative_umap` function.

Intra-point Euclidean distance was calculated for each space by comparing the spatial position of temporally adjacent points. Specifically, the position of a given point (corresponding to a unique temporal window) in behavior space (`x`_t_`, y`_t_) was compared to the position of the next point/window in time (`x`_t+1_`, y`_t+1_). The Euclidean distance between these points was then calculated and stored into a vector, allowing for the distribution of distances to be compared across replicates and parameter conditions. The coefficient of variation was also calculated for these measures to control for substantial variation in the magnitude of effect; this was done by dividing the distribution’s standard deviation by its mean.

Procrustes distance was used to measure the similarity between replicate behavior spaces of a certain parameter combination (Dryden & Mardia 1998). Briefly, this method treats spaces and their component points as a set of landmarks for pairwise comparison. To do so, pairs of behavior spaces are scaled to be similar sizes then shifted and rotated to possess the same position and orientation in space (Dryden & Mardia 1998). The distance between corresponding points in the two adjusted landmark sets is then calculated using Procrustes distance (reported here as Root Mean Square Error; RMSE). We performed this procedure and calculated Procrustes distance for all pairwise combinations of the 10 replicate behavior spaces within a given parameter set. Coefficient of variation for this measure was also calculated as above.

To assess recurrence, we measured the average return time for all points in each space. We first used the binning procedure outlined above to create a 32×32 grid so that the resulting space was composed of 1024 unique bins. We then computed a distance threshold to delineate a “neighborhood” around each bin (5% smallest intra-bin distance). For a given bin, each time the trajectory passed through its threshold the event was recorded and its duration stored. This distribution of return delays was analyzed by computing the proportion of points that displayed a return for a given time delay. For example, given a delay of 150ms, all bins would be scanned and the proportion possessing returns that occurred took between 150 and 160ms would be calculated. We calculated proportions in this manner for delays between 0ms and 2 seconds at 10ms intervals (as plotted in Figure 1I). Mean recurrence time was measured by collecting all observed return times for each replicate and calculating the mean.

### Generation of free-walking fruit fly behavior spaces

The movement trajectories of 20 female WTB flies walking in a circular area were collected from Haberkern et al. 2019. For each fly we computed velocity components as above and interpolated the resulting vectors from 12.3 Hz to 50 Hz to increase smoothness in the downstream behavior space. Initial analyses suggested the removal of 6 trials due to missing data or lack of substantial movement, resulting in a final dataset of 14 flies.

As above, we performed an empirical test to decide on the optimal window size for behavior space creation. To do so, we sampled the first 2000 frames from each trial (to increase computational efficiency) and swept through window sizes ranging from 20ms to 1 second (70 ms step size), creating a single behavior space for each of the 14 flies per window size. The resulting spaces were then directly compared using the Euclidean distance, Procrustes distance, and recurrence metrics previously described (Figure S2).

### Analyzing free-walking fruit fly behavior spaces

After sweeping window parameters, we generated a behavior space composed of all 14 free-walking trials using a window size of 160ms (399,869 windows). We then used `bin_space` to calculate point coordinates in a 64×64 grid. Stereotypy in movement through the space was visualized using a vector field transformation (Figure 2D). All instances in which trajectories through the space passed into a given bin were collected and then used to calculate the mean x and y vectors of the trajectory leaving the bin. These mean values were then represented visually using arrows that originated out of the corresponding bin, the direction and magnitude of which were dictated by the mean x and y vectors. Velocity distributions across the space (Figures. 2F-H) were visualized by calculating the mean value for each velocity component per window. These values were then used as input to the color function determining the hue of each point in the space.

Intra-fly variation in behavior space was assessed using the reduced 64×64 grid. For each fly we calculated the number of times its trajectory passed through a given bin, in addition to a binary measure of whether that bin was visited at all (‘0’ if not; ‘1’ if visited at least once). The continuous counts of bin visits were then used to calculate bin-wise mean and variance across all flies, the latter of which is visualized in Figure 3F. The binary measure was used to assess variation in overall space occupancy across flies (Figure S2).

We used 2-dimensional histograms to compare the overall behavioral patterns of individual flies. To do so, we created an individual map for each trial comparing the binned distributions of x and y coordinates via 2-dimensional kernel density estimation using the function `kde2d` (bandwidth = 2; 32 grid points in each direction) in the `MASS R` package. To facilitate comparisons across trials, the resulting density maps were normalized to the max density value of each and then linearized so that each was represented by a single vector of density values. The relationships between trials were inferred by calculating a distance matrix of these density vectors which was then used as input for hierarchical clustering (`hclust` function; Figure 3G).

### Generation of optogenetic local search behavior spaces

The movement trajectories of 19 Gr64f-Gal4 > ChrimsonR flies in virtual reality was collected from Haberkern et al. 2019. We processed and calculated velocity from these files as above, down sampling from 360 to 50 Hz to match the sampling rate of the free-walking dataset (for downstream comparisons). Windows were extracted from all 19 trials using the same width (160 ms) as above, yielding a library of 1,110,025 windows. We then used the `predict` function in the `UMAP R` package to embed all windows, one fly at a time, using the free-walking fly behavior space as a template. This procedure produced a combined behavior space composed of 33 trials (1,509,894 windows) that facilitated comparisons across experimental conditions and individuals.

Variation in space occupancy between free-walking and local search trials was assessed using 2-d density maps. As before, we used `bin_space` to get new coordinates for each trial, and then calculated density maps using `kde2d` (bandwidth = 1; 200 grid points in each direction), which were then normalized by dividing each density estimate by the maximum value in the map. As in Figure 3G these density maps were then linearized, combined in a matrix, and clustered using Euclidean distance and hierarchical clustering to produce the tree in Figure 4F.

### Analyzing optogenetic local search behavior spaces

We used a bin-wise Kruskal-Wallis test to statistically analyze differences in space occupancy between the two groups. For a given fly/trial we calculated the percent occupancy at each bin (number of visits to bin divided by total number of windows). A Kruskal-Wallis test was then used to compare the percent occupancies of the individual trials between free-walking and local search flies from which the test statistic and accompanying *p-*values were collected. We used Bonferroni correction to adjust these values, controlling for the number of tests performed (4,096). The adjusted *p-*values were then used to visually assess regions of greater differentiation between the free walking and local search trials, plotted as the −log10 transformation of the *p-*values (as seen in Figure 3F).

Time-evolving responses to optogenetic stimulation were assessed using density maps. To do so, we sampled the behavior space trajectories of each trial before and after all bouts of optogenetic stimulation. For each bout of stimulation, we extracted positions in behavior space for the second before stimulation in order to represent a baseline behavioral distribution. We then extracted positions immediately after stimulation using 1 second windows and step size of 100ms, from 0ms to 1 second afterward (Figure 4E). We combined positions across bouts and individuals within window to produce an aggregated response profile using kernel density estimation (bandwidth = 2; 64 grid points in each direction; common x and y limits across windows). If individuals displayed common behavioral responses to stimulation in a specific window, then the related density map should show structure in its distribution (i.e. concentrated red regions in Figure 4E).

### Larval Drosophila analysis

FIMtrack (Risse *et al*. 2014; Risse *et al*. 2017) was used to track the behavior of 72 *Drosophila* larvae while crawling on an agar surface (collected at 10Hz). FIMtrack outputs a number of per-frame measurements representation an animal’s shape and orientation. We selected primary measurements reflecting larval size, shape, and velocity (Figure 4A) in addition to the angular velocity of the head, midpoint, and tail for analysis. Due to variation in the mean of size measurements (i.e. area, perimeter, radii, spine length) over individual trials, these measures were detrended using the `ma` function in the `R` package `forecast` (window size = 10) after which all measures were then converted to z-scores. Given that the information of some of these features may be redundant, a principal component analysis was used to find an appropriate number of axes that could explain the variation in the dataset. We found that 8 principal components explained >90% of the variance in the feature set. These top 8 components were used as input to the iterative window procedure, sweeping a range of windows between 100ms and 5 seconds. As before the resulting spaces were compared using Euclidean distance, Procrustes distance, and recurrence metrics (Figures S3A-C). After analyzing these metrics, a window size of 800ms was chosen for all downstream analyses.

The full behavior space was plotted as vector field and with features highlighted in the fashion described above in the analyses of adult *Drosophila* behavior. Behavior labels and movement patterns through the behavior space (Figure 4D) were qualitatively assessed. Temporal patterns in the movement through behavior space were assessed via autocorrelation (Figure S3E). Autocorrelation was measured using the `acf` function in `R` (lag size = 100).

We used calcium imaging data from Tastekin *et al*. 2018 to examine neural activity during larval locomotion. We analyzed motor neuron activity (7 neurons per side) of larvae that were performing fictive locomotion (collected at 4-5Hz). Neural activity was measured using GcamP6f expressed in glutamatergic neurons (CG9887-lexA > GCamP6f) during optogenetic activation of PDM-DN neurons (PDM-DN>CsChrimson::mVenus) which induces stopping behavior. For each time point calcium fluorescence was converted to ΔF/F and then converted to z-scores for comparison across trials. A window size of 1 second was chosen for behavior space creation after the iterative window procedure. All trials were embedded in the same space and features/vector fields were plotted as previously described.

### Mouse 3D behavior analysis

Mouse 3D behavior data were previously published in Markowitz et al. 2018. In each trial, the mouse 3D posture was measured using the MoSeq pipeline (Wiltschko et al. 2015) from which the following features were calculated and used in this analysis: height, length, width, velocity (2-dimensional), velocity (3-dimensional), velocity (theta), and 10 postural principal components calculated from an 80 pixel x 80 pixel representation of the mouse’s position and height in 3D space. These features were then used an input to the iterative window procedure and a range of window sizes between 33ms and 1.66 seconds was examined (Figures S4A-C). A final window size of 133ms was chosen.

The full behavior space was plotted as vector field and with features highlighted as described above. Behavior labels and movement patterns through the behavior space (Figure 5D) were qualitatively assessed. The distributions of MoSeq syllables in behavior space (Figures 5K, S4E) were assessed by associating the timing of each syllable’s occurrence with the corresponding xy positions in behavior space. These were then used to calculate a 2-dimensional probability density function of the xy coordinates in space (100×100 grid). The density at each point in the 100×100 grid was then dividing by the maximum value so that the distribution varied between 0 and 1. The top 95% of these values were then plotted as a heatmap over the full behavior space distribution (Figures 5K, S4E).

The dispersion of syllables in behavior space was assessed via nearest neighbor distance using the function `nndist` (k = 1) in the `R` package `spatstat` (Baddeley et al. 2015). The significance of per-syllable dispersion was then computed via permutation tests. For each syllable, the mean nearest neighbor (nn) distance was calculated. The timing of syllable occurrence was then randomly shuffled 10,000 times. During each permutation the distribution of the shuffled data in behavior space was measured and used to calculate mean nn distance. Significance was assessed by computing *p-*values comparing the number of occurrences in which the shuffle nn distances were smaller than the observed mean nn distance, divided the number of permutations. Bonferroni correction was then used to adjust the *p*-values given the number of syllables tested.

A regularized generalized linear model was used to examine the relationship between mouse 3D pose behavior space and the original input features. The model was created with the `R` package `glmnet` (Friedman et al. 2010) using position in behavior space as the outcome variable and the features (height, length, width, velocity (2-dimensional), velocity (3-dimensional), velocity (theta), 10 PCs) as predictors. The set training set was composed of 75% of the data. We used 10-fold cross-validation via the `caret` (Kuhn 2016) package to compute the optimal alpha and lambda values for regularizing coefficient weights. The fit of the final model (r^2^) was computed by comparing the predicted behavior space positions in the remaining 25% of the data to the actual values. To measure variability across mice, individual models were also created for each trial in the same fashion, the fits of which are compared in Figure S4F.

### Generation of mouse 2D locomotor behavior space

Behavior data and neural recordings were previously published in from Hardcastle et al. 2017. For each cell, full trial spike event data (collected at 10kHz) and positional coordinates (collected at 50Hz) were extracted. As above, velocity components over the course of the whole trial were computed from the positional coordinates. We then used the empirical window-size test to identify the optimal window size for behavior space creation. 30 trials were randomly sampled for the iterative window test. The first 3000 frames of each trial were used to sweep through window sizes ranging from 20ms to 1 second (40ms step size; Figures S5A-F). The resulting spaces were then directly compared using the Euclidean distance, Procrustes distance, and recurrence metrics (Figures 6A-F). After analyzing these metrics a window size of 400ms was chosen for all downstream analyses.

Given the large size of the dataset (41,850,995 windows) we opted to perform a seed-embedding procedure to produce individual behavior spaces for each trial. To do so, we extracted and combined together the first 5,000 windows from each trial and created a seed behavior space using UMAP. We then individually embedded the remaining windows for each trial using the `predict` function in the `R` implementation of UMAP. Time points and XY coordinates corresponding to movement through these individual behavior spaces were then combined and used jointly for overall annotation of the mouse locomotion (As seen in Figures 6B-E). This all-trial behavior space was used to calculate 64×64 bin coordinates, allowing all mouse trials to be directly compared within the same behavior space architecture. Vector field representation (Figure 6D) and physical XY space movement decoding (Figure 6E) were computed as previously described.

The spike times for all cells in a given trial were then associated with the corresponding behavioral timepoints. To do so, spike rates were calculated over 20ms bins corresponding to the sampling rate of mouse positional data (50Hz) and, thus, the rate of movement through behavior space. The rate of activity was also calculated over 1 second (1Hz) scales to explore the extent to which broad temporal differences might be present between neural activity and behavior. For each cell/trial pair the Pearson correlation between behavior space position and firing rate was calculated. The significance of these correlations was measured using permutation tests. For each cell, the spike rate data were shuffled 10,000 times and then correlated with position in behavior space. These correlations were used to calculate *p-*values by comparing the number of times shuffled correlations were greater than the observed value, divided by the number of permutations (10,000; Figure 5I). 2-dimendionsal tuning curves (as in Figures 6F-G and Figure S5M) were computed for each cell by calculating the mean spike rate in each bin using the 64×64 representation previously calculated. Mutual information between the cells and behavior space was calculated (in bits) by comparing the resulting 2-d tuning curves to the distribution of average occupancy time per bin for each trial using the function `mi.plugin` from the `R` package `entropy`. The canonical MEC coding variables (speed score, spatial stability, head direction, border score; Figures 6K-R) used were previously computed following Hardcastle *et al*. 2017. Cross-correlations between neural activity and behavior space position were calculated with the function `ccf` in `R` (lag of 16 seconds as seen in Figure 7C).

Temporal variation in the association of neural activity and behavior space (as in Figures 7D-H) was assessed by calculating the correlation coefficient, entropy, and mutual information between the two across a range of windows (0-4 seconds, 20ms steps). Correlations and mutual information were calculated for each window size as above. Shannon entropy (in bits) was calculated using the distribution of mean firing values from 2-d neural tuning curve for each window (using the `entropy` function from the `entropy R` package). MI maximized window sizes were chosen for each cell using the timescale at which the maximum mutual information between behavior space position and neural activity occurred.

Rate-based differences in coding were assessed using the MI maximized window sizes identifying above. For each cell, we calculated the activity rates corresponding to the 75th, 80th, 90th, 95th, and 99^th^ percentiles. We then identified the time points at which each cell’s activity occurred above the respective percentile. The corresponding positions in behavior space were then extracted in 8 second windows surrounding each timepoint (from 4 seconds before to 4 seconds after; 20ms bins). 2-d tuning curves were then constructed from the distribution of points in behavior space for all events that occurred across the corresponding 20ms bins. For example, the behavior space positions occurring 100-80ms before all events above the 99^th^ percentile would be compared, followed by the same calculation for all events occurring 80-60ms before, then 60-40ms before, etc. This procedure would thus produce 400 2-d tuning curves per percentile, corresponding to a sampling rate of 50Hz over a course of 8 seconds. The Shannon entropy (in bits) of each 2-d tuning curve was then calculated and plotted as a curve over time to detected consistent changes in its distribution (Figure 7K). To compare these entropy distributions across all cells each was divided by its maximum value. The normalized entropy measures for the 75^th^ and 99^th^ percentile values were then used as input to hierarchically cluster all cells. Clusters were identified using a dynamic tree cutting algorithm via the `cutreeHybrid` function in the R package `dynamicTreeCut` (minimum cluster size of 10) (Langfelder et al. 2007).

---

**Figure S1.**
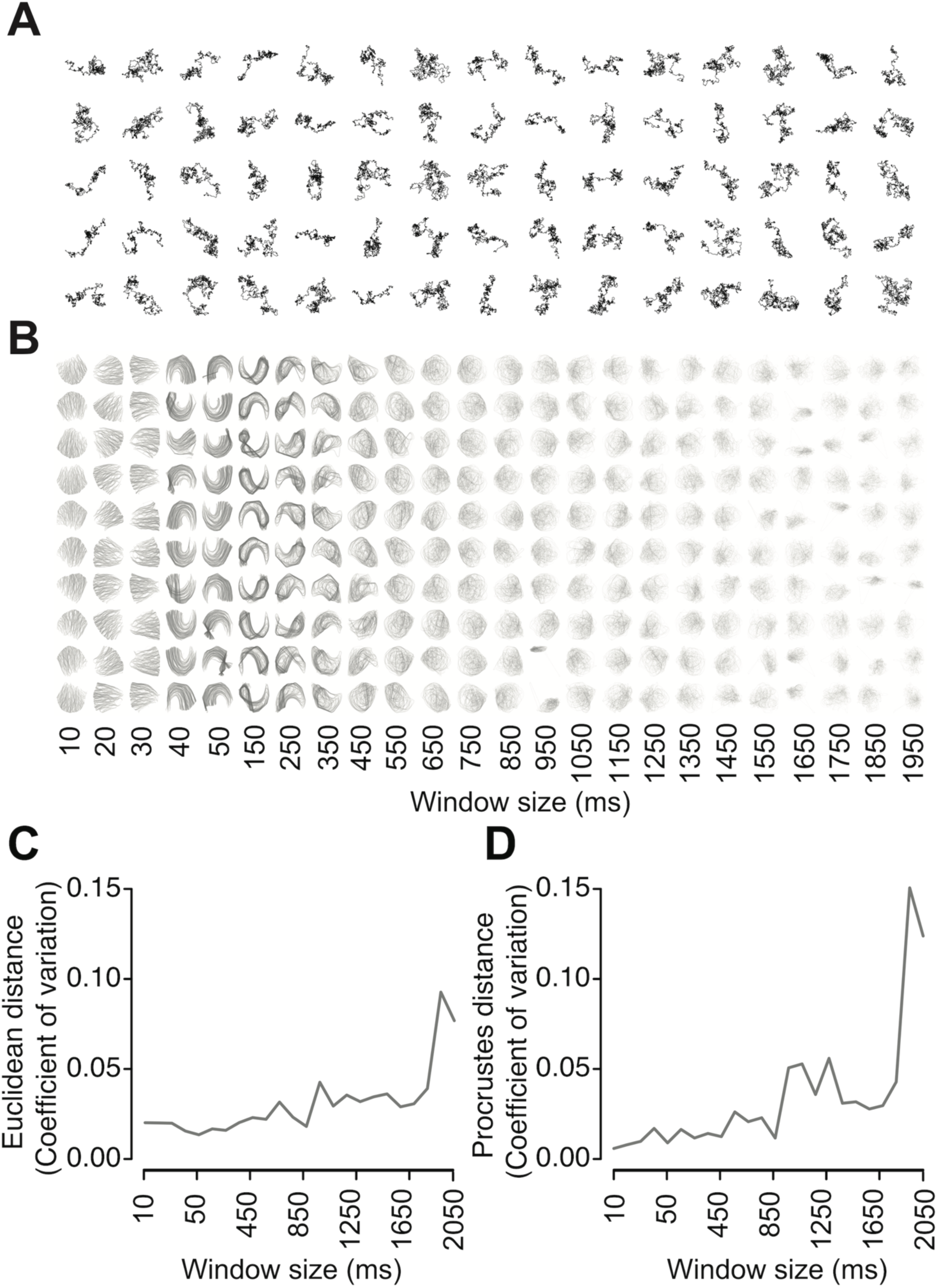
Correlated random walk iterative window tests. **(A)** A set of 100 example correlated random walks used for parameter tuning. Each is 10,000 frames long and sampled at 100 FPS. **(B)** Trajectories through behavior space are plotted for each of the 10 correlated random walk replicates per window size tested. Replicates are arranged in rows while window sizes are arranged in columns. Trajectories were visualized by connecting temporally adjacent points in behavior space with lines. Recurrent dynamics are represented by highly overlapping lines, reflecting repeated excursions through those specific paths. **(C)** Coefficient of variation of mean intra-point Euclidean distance as a function of window size. **(D)** Coefficient of variation of Procrustes distance RMSE as a function of window size.

**Figure S2.**
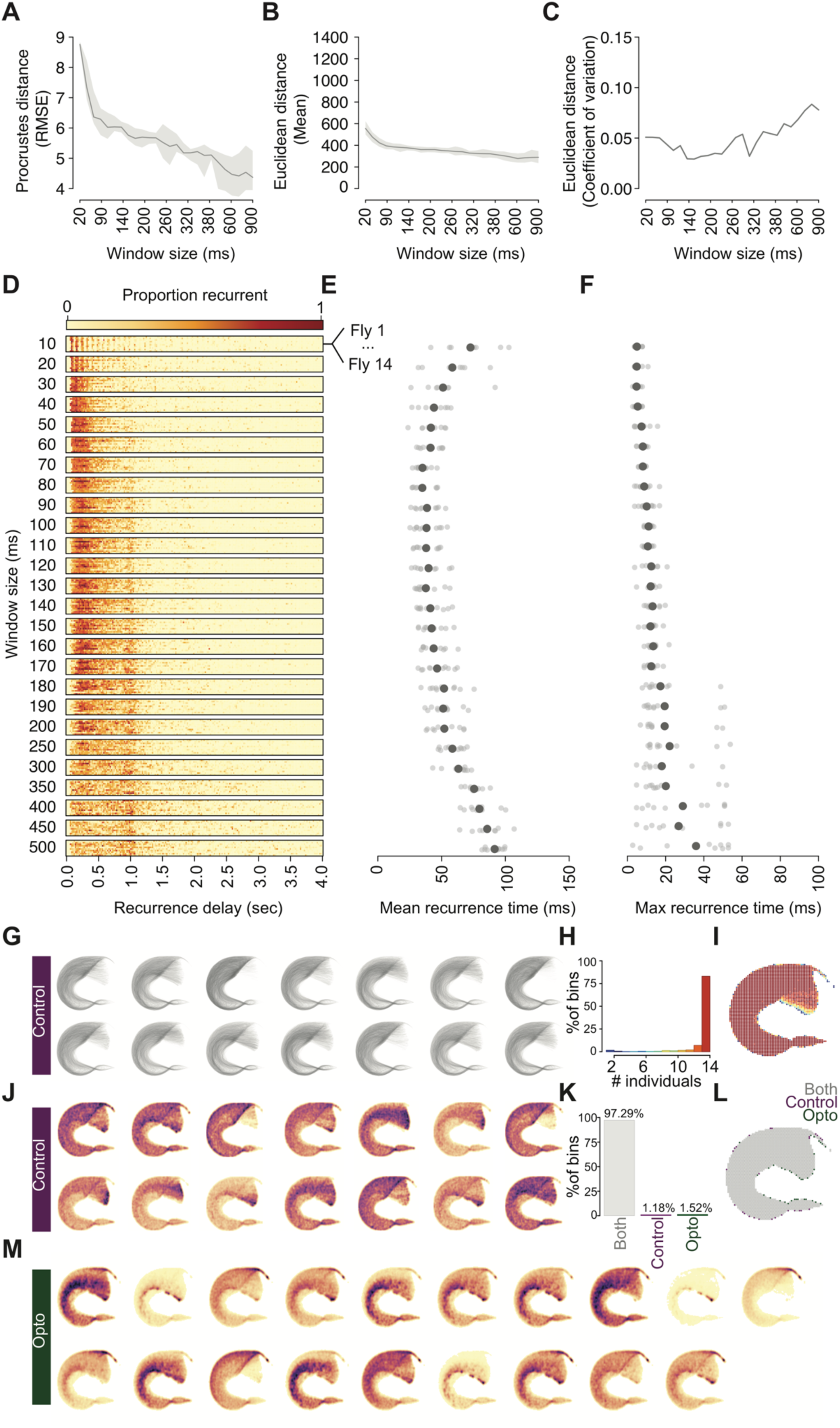
Assessing structural and temporal components of the free-walking fly dataset. **(A)** Procrustes distance RMSE measures as a function of window size. The darker line corresponds to the mean value while the shading reflects standard error. **(B)** Mean intra-point Euclidean distances. The darker line corresponds to the mean value while the shading reflects standard error. **(C)** Coefficient of variation of mean intra-point Euclidean distance as a function of window size. **(D)** Recurrence plot of behavior spaces produced from the free-walking dataset. The proportion of recurrent points given a range of time delays ranging from 0 to 4 seconds is indicated by the color of the corresponding bins (from light yellow to dark red). Each horizontal window size bar includes all 14 flies tested. **(E)** Mean recurrence times for all 14 flies as a function of window size. The larger circle corresponds to the population mean while smaller circles correspond to each fly. **(F)** Maximum recurrence times for all 14 flies as a function of window size. Each point reflects the bin in which the largest proportion of points displayed recurrence. The larger circle corresponds to the population mean while smaller circles correspond to each fly. **(G)** Pathways through the locomotor behavior space for all 14 control flies, produced by connecting temporally adjacent windows with partially transparent lines. **(H)** Bar plot comparing the percent of behavior space bins that were visited by at least *n* individuals. Bars are color coded from blue to red to correspond to **(C)**. **(I)** Locomotor behavior space binned to a 64×64 grid. Each bin is colored corresponding to the number of individuals that visited it. **(J)** Density maps for all control flies. Each behavior space corresponds to an individual and is colored by the kernel density estimate generated for that fly’s trajectory in behavior space. Darker color corresponds to a greater density in a given region. **(K)** Bar plot comparing the percentage of bins in a 64×64 gridded behavior space that were visited by control flies (purple), optogenetically activated flies (green), or both (grey). **(L)** Behavior space colored by the distribution of overlapping and unique bins. Each bin is colored by the designations in the bar plot. **(M)** Density maps for optogenetically activated flies. Each behavior space corresponds to a unique fly (n = 19) and is colored by the kernel density estimate generated for that fly’s trajectory through behavior space. Darker colors correspond to a greater density in a given region.

**Figure S3.**
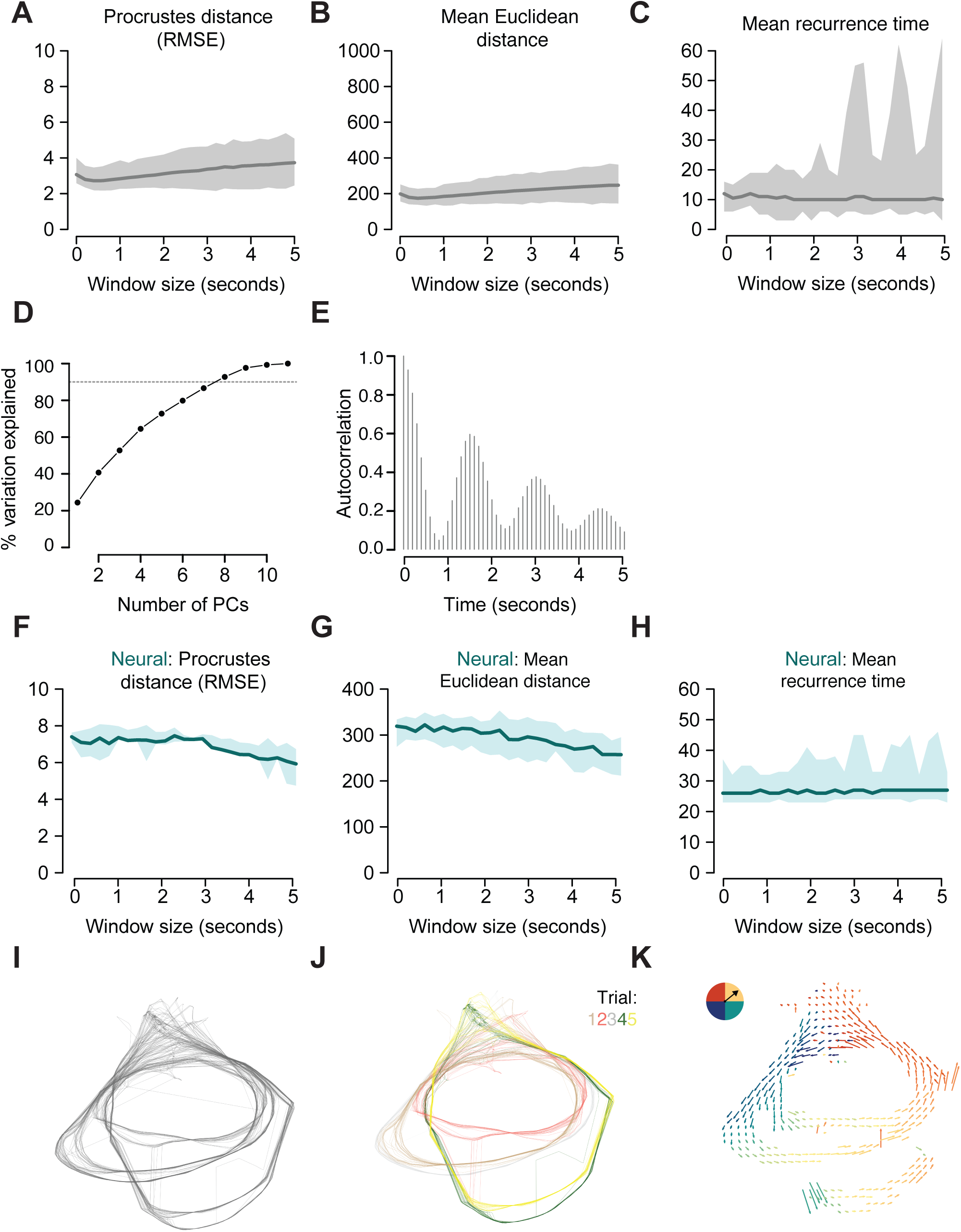
*Drosophila* larvae behavioral and neural spaces. **(A-C)** Procrustes distance **(A)**, mean Euclidean distance **(B)**, and mean recurrence time **(C)** as a function of window size for the crawling *Drosophila* larvae behavior space. The darker line corresponds to the mean value while the shading reflects standard error. **(D)** Scatterplot of the percent variation explained by increasing numbers of principal components representing the 11 input features used. Dashed line corresponds to 90% variation explained. **(E)** Example distribution of the autocorrelation of position in larval behavior space over 200 seconds. Position was represented by single value encoding of the 64×64 binned behavior larval space and then used as input to the autocorrelation calculation. **(F-H)** Procrustes distance **(F)**, mean Euclidean distance **(BG**, and mean recurrence time **(H)** as a function of window size for the crawling *Drosophila* larvae neural space. In this case, the iterative windows procedure was run using fluorescence traces from 7 motor neurons as input. The darker line corresponds to the mean value while the shading reflects standard error. **(I)** The full larval *Drosophila* neural space, containing 5 independent trials. **(J)** The full larval *Drosophila* neural space, colored by individual trials (denoted in legend in upper right hand corner). **(K)** Larval *Drosophila* neural space represented as a mean vector field. Color corresponds to the angle of each vector (reflected by colored circle in the upper left hand corner).

**Figure S4.**
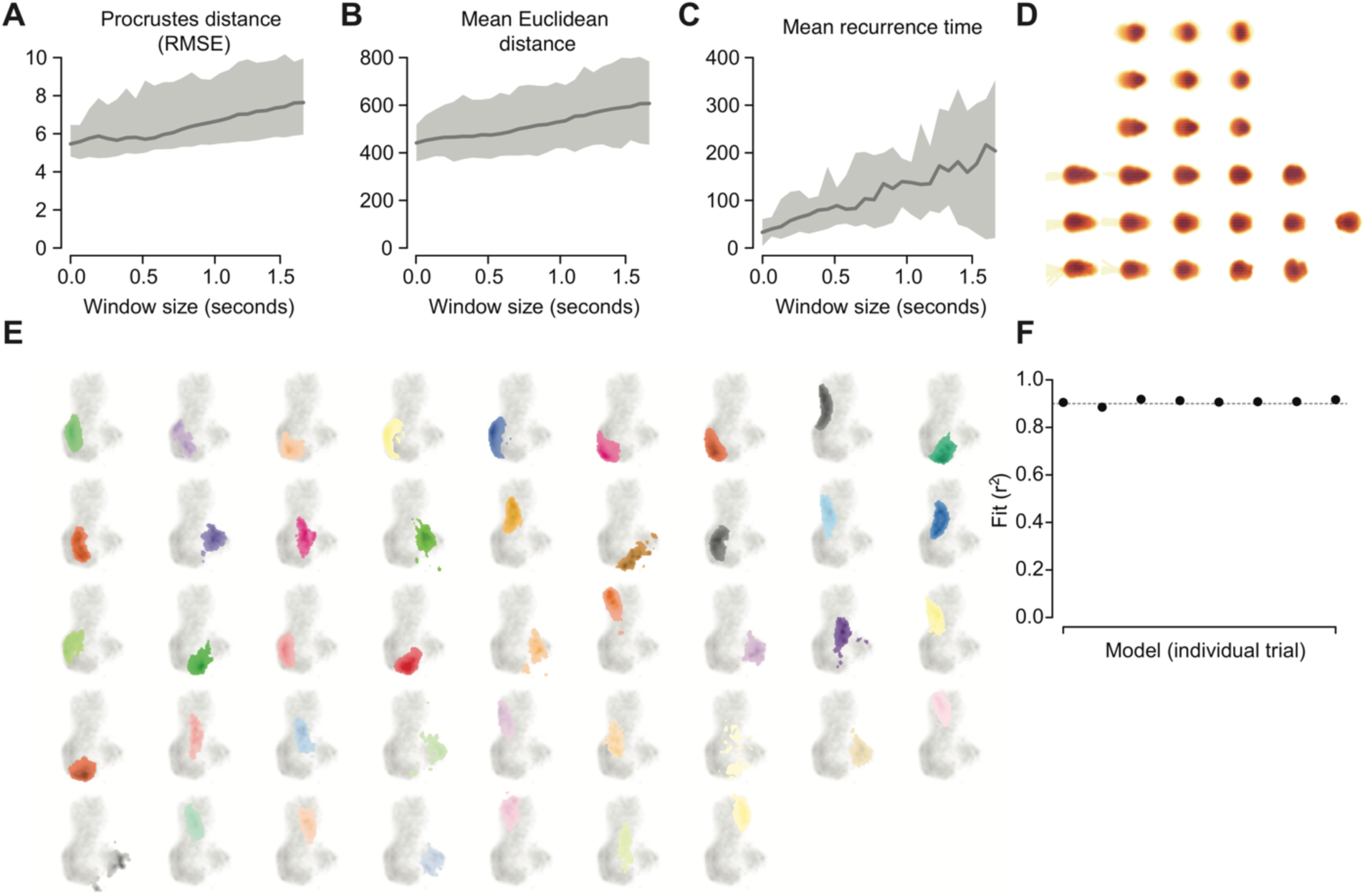
Mouse 3D pose behavior space. **(A-C)** Procrustes distance **(A)**, mean Euclidean distance **(B)**, and mean recurrence time **(C)** as a function of window size. The darker line corresponds to the mean value while the shading reflects standard error. **(D)** Heatmaps representing mean 3D pose as a function of behavior space position (grouped into 25 unique bins). Time points in which each bin was visited were extracted and then associated with the corresponding moments in the raw 3D imaging data. These instances were then averaged, producing a mean 3D posture per bin, represented here as a heatmap (yellow = further from imaging camera; darker red = closer/higher). **(E)** The distributions of the full set of behavioral syllables (identified by MoSeq) in behavior space. A probability density function across behavior space was computed for each syllable and then plotted in color on top of the full behavior space (in grey; as in Figure 5K). **(F)** Fits (r^2^) of generalized linear models comparing input features and behavior space position for individual trials/mice. The fit of the model using all trials is denoted by the grey dashed line.

**Figure S5.**
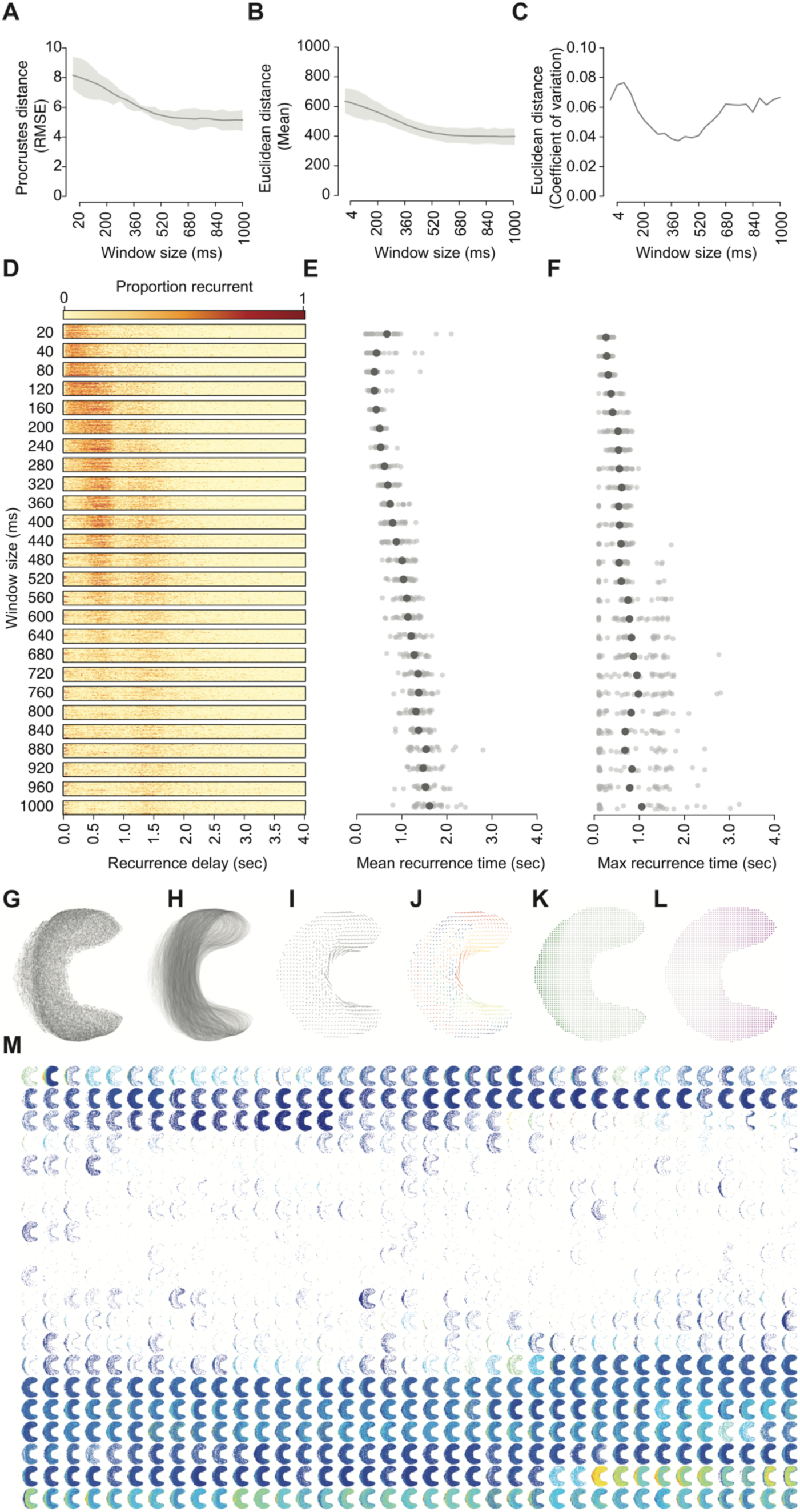
Mouse locomotor behavior space and its association with MEC activity. **(A)** Procrustes distance RMSE measures as a function of window size. The darker line corresponds to the mean value while the shading reflects standard error. **(B)** Mean intra-point Euclidean distances. The darker line corresponds to the mean value while the shading reflects standard error. **(C)** Coefficient of variation of mean intra-point Euclidean distance as a function of window size. **(D)** Recurrence plot of behavior spaces taken from 30 random mouse trials. The proportion of recurrent points given a range of time delays ranging from 0 to 4 seconds is indicated by the color of the corresponding bins (from light yellow to dark red). Each horizontal window size bar includes all 30 mice tested. **(E)** Mean recurrence times as a function of window size. The larger circle corresponds to the population mean while smaller circles correspond to each mouse trial. **(F)** Maximum recurrence times as a function of window size. Each point reflects the bin in which the largest proportion of points displayed recurrence. The larger circle corresponds to the population mean while smaller circles correspond to each mouse trial. **(G)** Mouse locomotor behavior space, each point corresponds to a temporal window. **(H)** Pathways through mouse locomotor behavior space, produced by connecting temporally adjacent windows with partially transparent lines. **(I)** Mouse locomotor behavior space represented as a vector field. Arrow direction and magnitude correspond to the angle and mean direction taken after visiting each bin. **(J)** Arrow direction and magnitude correspond to the angle and mean direction taken after visiting each bin. Arrows are colored by the degree of the direction vector (corresponding to circle in upper left-hand corner). **(K)** Distribution of translational velocity across mouse behavior space (darker green corresponds to higher values). **(L)** Distribution of angular velocity across mouse behavior space (darker purple corresponds to higher values). **(M)** 2-d tuning curves of neural activity (1 second bins) for all MEC neurons. The ordering of tuning curve position was determined by hierarchical clustering (i.e. more similar tuning curves are placed adjacent to each other).

**Figure S6:**
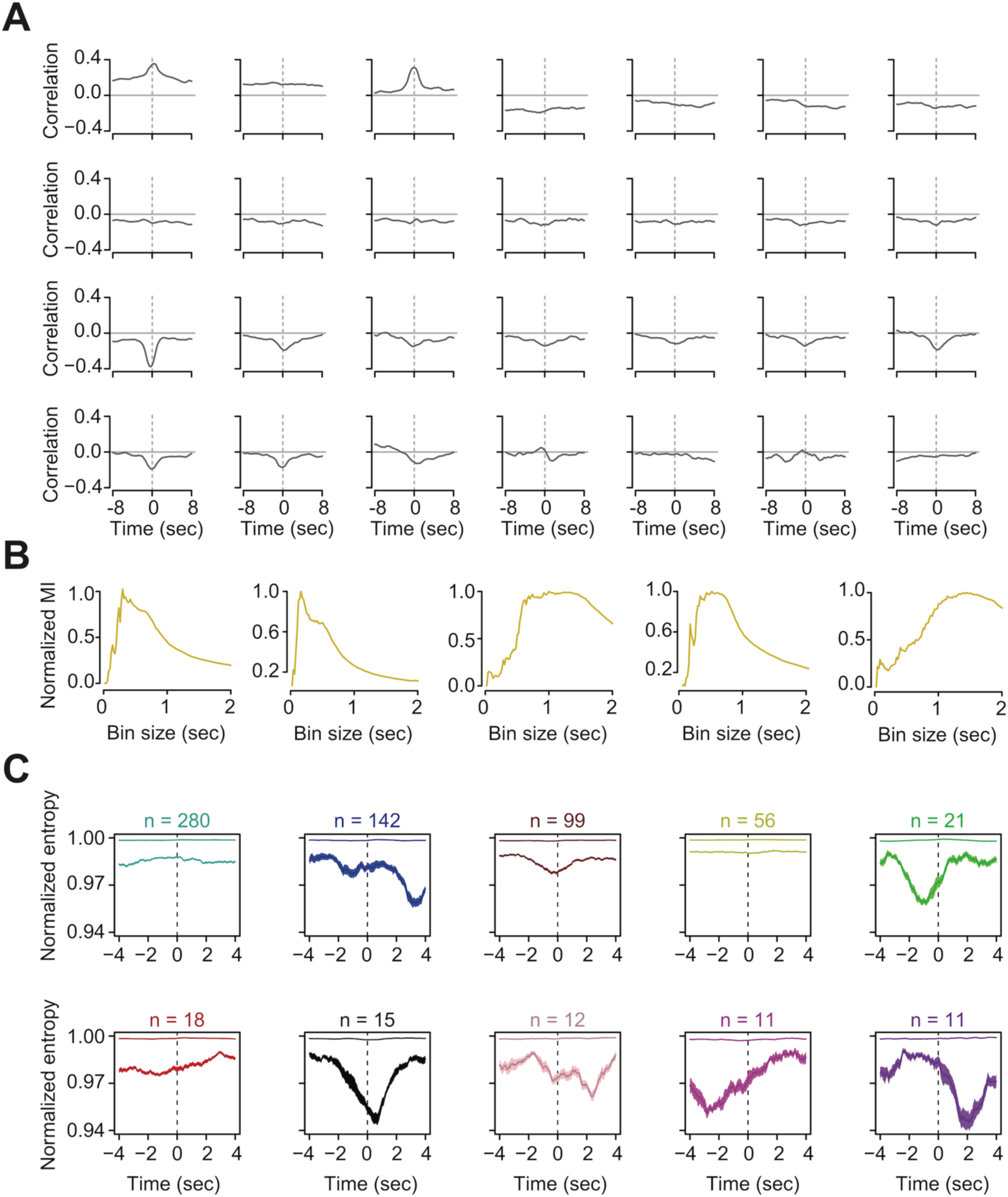
Variation in temporal and rate-based associations between MEC neurons and behavior. **(A)** Example cross-correlations between behavior space position and MEC neuron activity over a 16 second time window. The dotted grey line corresponds to time point zero. The solid grey line reflects a correlation coefficient of zero. **(B)** Example mutual information (MI) tuning curves as a function of binning size of neural activity. For comparison purposes, MI is presented as a normalized value in which each distribution has been divided by its maximum value. **(C)** Mean entropy distributions of ten clusters identified using dynamic tree trimming. Plotted are the distributions for the 75% (top) and 99% (bottom) cutoffs. Mean and standard error are plotted as well as sample sizes, as in Figure 7E.

